# Early Disruption of Photoreceptor Cell Architecture and Loss of Vision in a Humanized Pig Model of Usher Syndrome

**DOI:** 10.1101/2021.05.31.446123

**Authors:** Sophia Grotz, Jessica Schäfer, Kirsten A. Wunderlich, Zdenka Ellederova, Hannah Auch, Andrea Bähr, Petra Runa-Vochozkova, Janet Plutniok, Vanessa Arnold, Taras Ardan, Miroslav Veith, Gianluca Santamaria, Georg Dhom, Wolfgang Hitzl, Barbara Kessler, Mayuko Kurome, Valeri Zakharchenko, Joshua Linnert, Andrea Fischer, Andreas Blutke, Anna Döring, Stepanka Suchankova, Jiri Popelar, Helen May-Simera, Karl-Ludwig Laugwitz, Luk H. Vandenberghe, Eckhard Wolf, Kerstin Nagel-Wolfrum, Jan Motlik, M. Dominik Fischer, Uwe Wolfrum, Nikolai Klymiuk

**Author notes:** contributed equally to the study, shared 1st author. contributed equally to the study, shared last authorship (Uwe Wolfrum, Nikolai Klymiuk). KAW present address: Department of Physiological Genomics, LMU Munich, Germany. Correspondence should be addressed to N.K. and U.W.

## Abstract

Usher syndrome (USH) is the most common form of monogenic deaf-blindness. Loss of vision is untreatable and, so far, there are no suitable animal models for testing therapeutic strategies. By introducing a human mutation into the harmonin-encoding *USH1C* gene in pigs, we generated the first translational animal model for USH type 1 with characteristic hearing defect, vestibular dysfunction and visual impairment. Changes in photoreceptor architecture, quantitative motion analysis and electroretinography were characteristics of the reduced retinal virtue in USH1C pigs. Primary cells from those animals and USH1C patients showed significantly elongated primary cilia, compared to wild-type, confirming the nature of USH as a true and general ciliopathy and proving the therapeutic capacity of gene supplementation and gene repair approaches.

## Introduction

Usher syndrome (USH) is the most common form of inherited deaf-blindness in humans (*1*). USH is clinically and genetically heterogeneous, with at least 12 genes assigned to three clinical USH types (*2, 3*). The most severe of them is USH1, characterized by profound hearing loss from birth on, vestibular areflexia, and pre-pubertal onset of Retinitis pigmentosa (RP). In patients, the congenital sensorineural hearing impairment can be compensated with cochlear implants, but no therapeutic option is presently available for the ocular disease component. Thus, most USH cases lead to severe visual impairment and blindness over time, adding a substantial psychological component to the clinical symptoms (*4*).

Development of therapies and understanding of USH pathogenesis in the retina is arduous as existing rodent models for USH reflect the human deficits in the inner ear, but show only a very mild, if any, retinal phenotype (*5, 6*). This has been correlated to substantial differences between mouse and human in anatomy and cellular composition of the retina and stimulated discussion about alternative model species. Pigs reflect structure and function of the human eye much better than mice as they have a similar size, a comparable rod/cone ratio and a cone-rich region, the so-called visual streak (*7, 8*). Specifically, calyceal processes (CP), cellular extravaginations at the transition of the inner to the outer segment (OS) of photorepetor cells (PRC) that comprise USH proteins (*9, 10*) and appear in many species, including human and pig, but not rodents (*5, 11, 12*). Various pig models have been described for retinal degeneration, with the majority of them induced by compounds or light (*13–15*) or additive transgenesis (*16–19*). As the latter do not truly reflect the genetic constellation in patients, the phenotype was, however, difficult to interpret and limited the utility of these models.

Here, we describe a pig model for USH1 (USH1C pig), that obliterate harmonin, a major scaffold protein in the USH interactome (*20, 21*) upon humanizing a fragment carrying a patient-specific mutation in the *USH1C* gene. In addition to profound hearing impairment and pronounced vestibular dysfunction, USH1C pigs revealed a significant visual dysfunction as well as impaired ciliogenesis in primary cells, facilitating the evaluation of gene complementation therapy or human-specific gene repair.

## Results

### A patient-specific disease-causing segment in *USH1C* leads to Usher syndrome in pigs

First, we assessed the translational potential of an USH pig model by examining the harmonin proteins and the underlying genetic *USH1C* structure across mammalian species. With the exception of a few small segments in the coding regions, proteins show an outstandingly high degree of conservation between the species (Figure 1a). Extended CTCF/cohesin-binding regions suggest that exons undergoing complex splicing are controlled by distantly located chromosomal segments in *cis* rather than by local regulatory elements (Figure 1b). In contrast, a 1.5kb segment comprising exon 2 and its surrounding intronic regions were predicted to be relevant in gene regulation (Figure 1c). Conservation is noticeable throughout the entire region, culminating in a 100% identity among all amino acid positions encoded by exon 2 in all examined species (Figure 1d). We therefore modified the porcine *USH1C* gene by a corresponding 1.5kb element including a disruptive C91T/R31X nonsense mutation (*22*) as well as 3 intronic SNPs identified on the disease-allele of a human USH1C patient.

**Figure 1.**
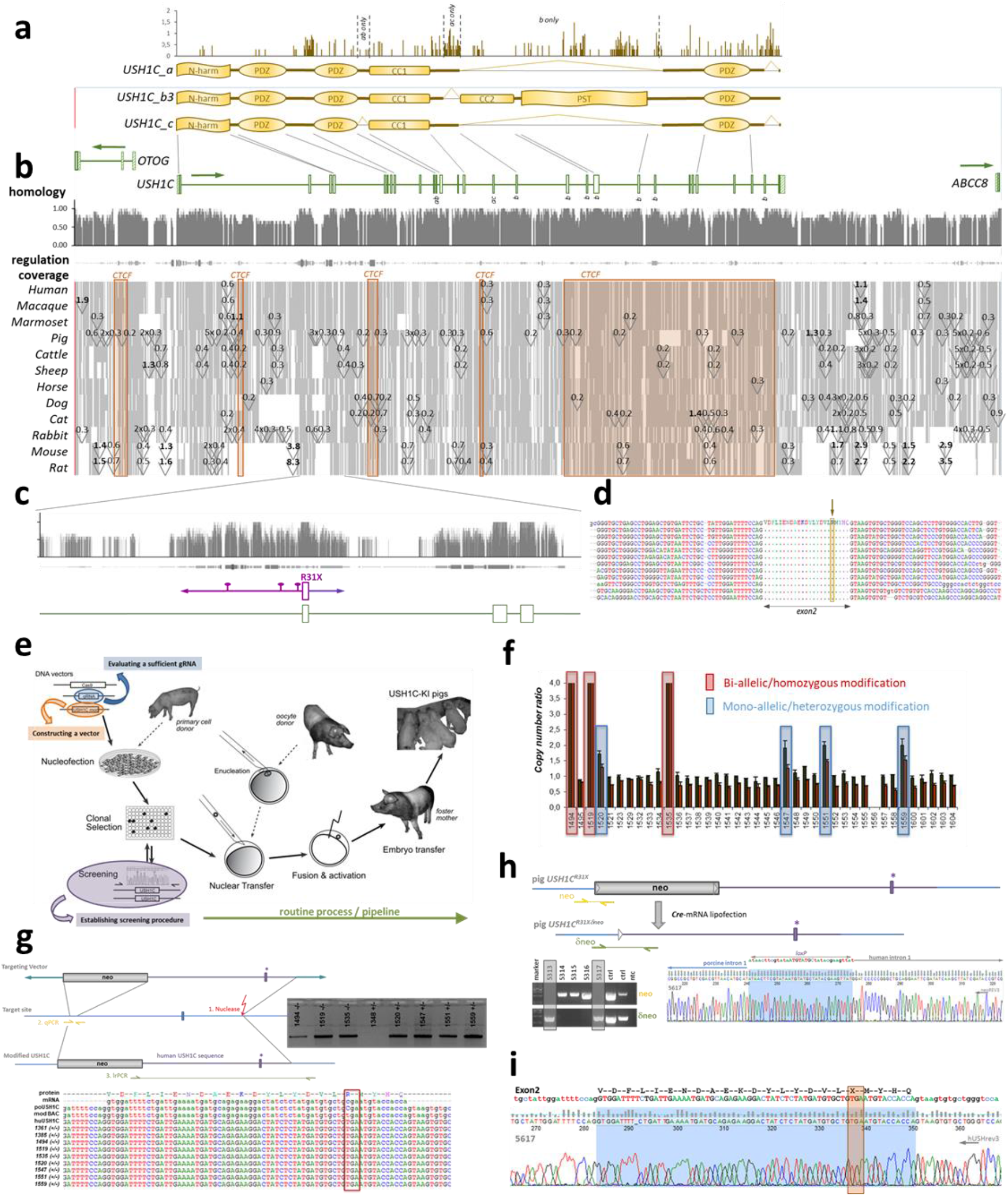
Generation of USH^R31X^ pigs. **(a)** Three main splice variants a, b3 and c have been confirmed for human *USH1C*. According to an Entropy plot (upper panel), the coding regions appear highly conserved among mammalian species with some variability in inter-domain segments, but also within the b3-specific PST domain. A high degree of conservation occurs also at nucleotide level (**b**, upper panel), including exons and intronic elements that appear relevant for transcriptional regulation (middle panel). Most of the *USH1C* gene and its surrounding regions are consistently abundant in the examined species (lower panel) and only few intronic areas are segmented by multiple repetitive elements (triangles, with approximate length in kb). Indicated as CTCF regions, significant proportions of the genes seems to be involved in chromosomal interactions (orange blocks). The 5′-end of USH1C is characterized by a high degree of sequence conservation at nucleotide level and strong evidence for regulatory purposes **(c)**, the lack of repetitive elements and identity at protein level **(d)**, suggesting the humanization of a consistent region around exon 2 (magenta in c). Based on determination of patient-specific sequencing, the modification contained the causative position for the R31X mutation (arrow in d) and 3 additional intronic patient specific SNPs (pins in c). **(e)** USH1C^R31X^ founder animals were established by cloning pig primary cells that have been nucleofected with plasmids encoding the unspecific Cas9, gRNA4 (blue) and a modified BAC as targeting vector (orange). Single cell clones were generated and characterized (magenta) by a qPCR-based LOWA assay **(f)** and highlighted clones with mono-allelic (blue) or bi-allelic (red) modifications. Introduction of the humanized fragment was confirmed by Sanger-sequencing of a long-range PCR **(g)** and verified the abundance of the TGA non-sense codon at position 31 (red box). Before entering the SCNT pipeline, single cells were treated with Cre-mRNA and the delivered piglets were examined for the excision of the selection cassette **(h)**. Finally, founder animals were examined for the desired modification at genomic level, including the non-sense codon (orange box) in exon 2 (blue) **(i)**.

Following previous attempts to generate pig models (*23, 24*), somatic cell nuclear transfer (SCNT) was used for establishing founder animals from genetically modified pig primary cells (Figure 1e). Into those cells, the 1.5kb human fragment as well as a floxed selection cassette were introduced by combining a BAC vector with CRISPR/Cas components. (Figure S1a-f). Upon clonal selection, primary cells revealed mono- or bi-allelic modification of the *USH1C* locus (Figure 1f, g). Female cell clones with both *USH1C* alleles modified were further lipofected with Cre-mRNA in order to excise the selection cassette prior to their usage in SCNT experiments (Figure 1h). Out of 12 embryo transfers, 4 litters were produced, delivering a total of 18 piglets. Genetic analysis confirmed the presence of the humanized fragment as well as correct transitions from the construct to the porcine sequence in the animals (Figure 1i). Excision of the *neo* cassette was achieved in 5 out of 15 founder animals, leaving a single lox site remaining (Figure 1h).

Strikingly, all cloned founder animals showed a pronounced circling phenotype at birth (Video S1), correlating with a dysfunctional vestibular system (*25, 26*). For this reason, animals were raised in a rescue deck from birth on (*27*). After 48-72 h of interval feeding by hand, circling essentially ceased and USH1C piglets were able to feed themselves from the nurturing unit.

In general, USH1C pigs developed and acted normally, but stress-induced circling proved a consistent hallmark in the case animals found themselves challenged by new situations. At an age of 3-6 months, we occasionally observed nystagmus (Video S2), a common phenomenon of vestibular dysfunction and vertigo (*28*) and previously described in USH patients (*29–31*). 8-months USH1C pigs were fertile and 3 sows became pregnant after insemination and gave birth to F1 litters, comprising heterozygously affected offspring. F2 USH1C piglets were produced by mating F1 animals with each other or by inseminating the cloned founder sows with sperm of one of their heterozygous offspring. In all USH1C offspring, a circling phenotype appeared as it had been observed in the F0 generation produced by SCNT.

### Motion analysis demonstrates vestibular dysfunction and visual deficits in USH1C pigs

To quantify changes in visually guided behaviour, animals were challenged in distinct settings. In a barrier course in which animals had to bypass vertical shields to reach a food bowl (*32*), USH1C pigs did not show significant differences to WT controls regarding duration or gait instability (Videos S3, S4 and Figures 2a, S2). Interestingly, however, USH1C animals hesitated to enter the course in the dark (average 2.9 lux), turned around more often within the course and had more frontal contacts with obstacles than WT control animals. In a more complex obstacle course, in which animals had to cross or bypass distinctly shaped hurdles (Videos S5, S6 and Figure S3), USH1C pigs had significantly more contacts with hurdles while their speed was similar to WT in the dark (0.1 - 12 lux) (Figures 2b, S4). In the light (50-150 lux), USH1C pigs did not only touch hurdles more often, but were also slower than WT controls. Remarkably, USH1C pigs had difficulties to pass hurdles taking a pronounced vertical movement and therefore requiring balance (such as the red cavaletti) as well as obstacles challenging stereopsis and peripheral vision (such as the hanging or standing blue barrel and the sideboard) (Figure 2c). While the interpretation of an impaired vestibular system was further supported by the clearly different gait in USH1C pigs (Videos S5, S6), the interpretation of a reduced visual function in these animals was corroborated by a more detailed analysis of the movement in the obstacle course (Figure 2d). It became obvious that USH1C pig trajectories are longer (TrajLength and TrajDistance) compared to WT, and they often change direction abruptly (lower Emax). In addition, they take more time to complete the course (TrajDuration) and their step length is shorter.

**Figure 2.**
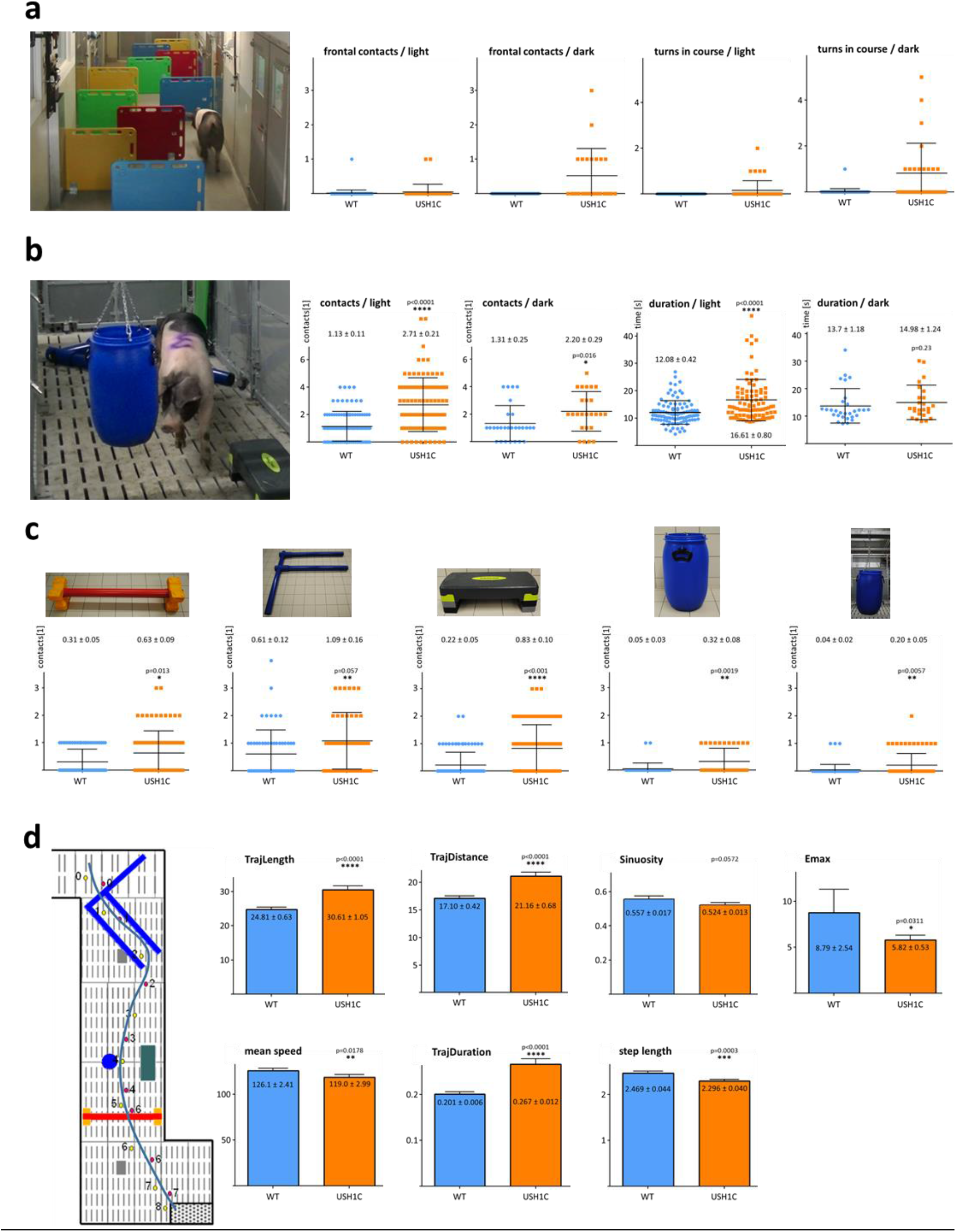
Impaired movement of USH pigs in 3-dimensional space. In barrier course **(a)**, different arrangements of shields *(see Figure S2)* were used to challenge orientation. Distinct parameters were examined, either in the dark or under normal light conditions and revealed only minor differences between WT (blue) and USH1C (orange) pigs. Mean values ± standard deviation are given by lines. **(b)** In a similar setting, different obstacles were placed instead of shields *(see Figure S3)* and revealed intensified sensing by snout contacts for USH1C pigs in the dark and in under normal light. In the latter case, WT pigs were also faster under normal light, whereas speed was similar in the dark. Mean values ± standard deviation are given by lines, mean values ± standard error are given by numbers, p-values for Mann-Whitney U-tests are indicated. Analysis included all runs, for normalized data sets, see Figure S4. **(c)** Detached analysis revealed that both, obstacles primarily challenging the vestibular system (red cavalletty, blue F) as well as hurdles daring vision (sideboard, hanging or standing ton) provoked significant differences between WT and USH1C pigs. **(d)** Plotting individual runs to a planar matrix facilitated parametrizing the positions of the left (red) and right front leg (red) as well as the time when animals touch the ground (in seconds) and comparison to the positions of the hurdles. Data were then introduced in TrajR for evaluating standard motion parameters. Coordinates in the parcour were indicated in cm and Y-axis of the respective parameters are given according the TrajR output.

### Clinical examinations confirm hearing loss and vision impairment in USH1C pigs

Auditory brainstem response (ABR) documented severe hearing deficits in USH1C pigs as early as 3 weeks and up to 2 years of age when response was lacking even upon a click stimulus at sound pressure levels of up to 120dB SPL (Figures 3a, S5). The findings of sensoneural hearing loss(*33*) correlated with changes in the arrangement of hair bundles and lacking hair cell stereocilia in the inner ear (Figure S5b), the findings in *Ush1c* mouse models (*26, 34*) and the congential deafness in human patients (*35*). The hearing threshold of heterozygous USH1C^+/−^ pigs was slightly elevated compared to WT littermates (Figure S5e, f) and for both genotypes it decreased with frequency while it increased in USH1C animals (Figures 3a). In line with that, USH1C piglets did not react on their mother’s call for nourishing (Suppl_Video 7). Most of them needed nudging for waking up, while heterozygous littermates arrived at their mother within 15 seconds (Figure 3a). Consistently, distinct USH1C piglets were within the faster group, presumably awoken from being sideswiped by their littermates.

**Figure 3.**
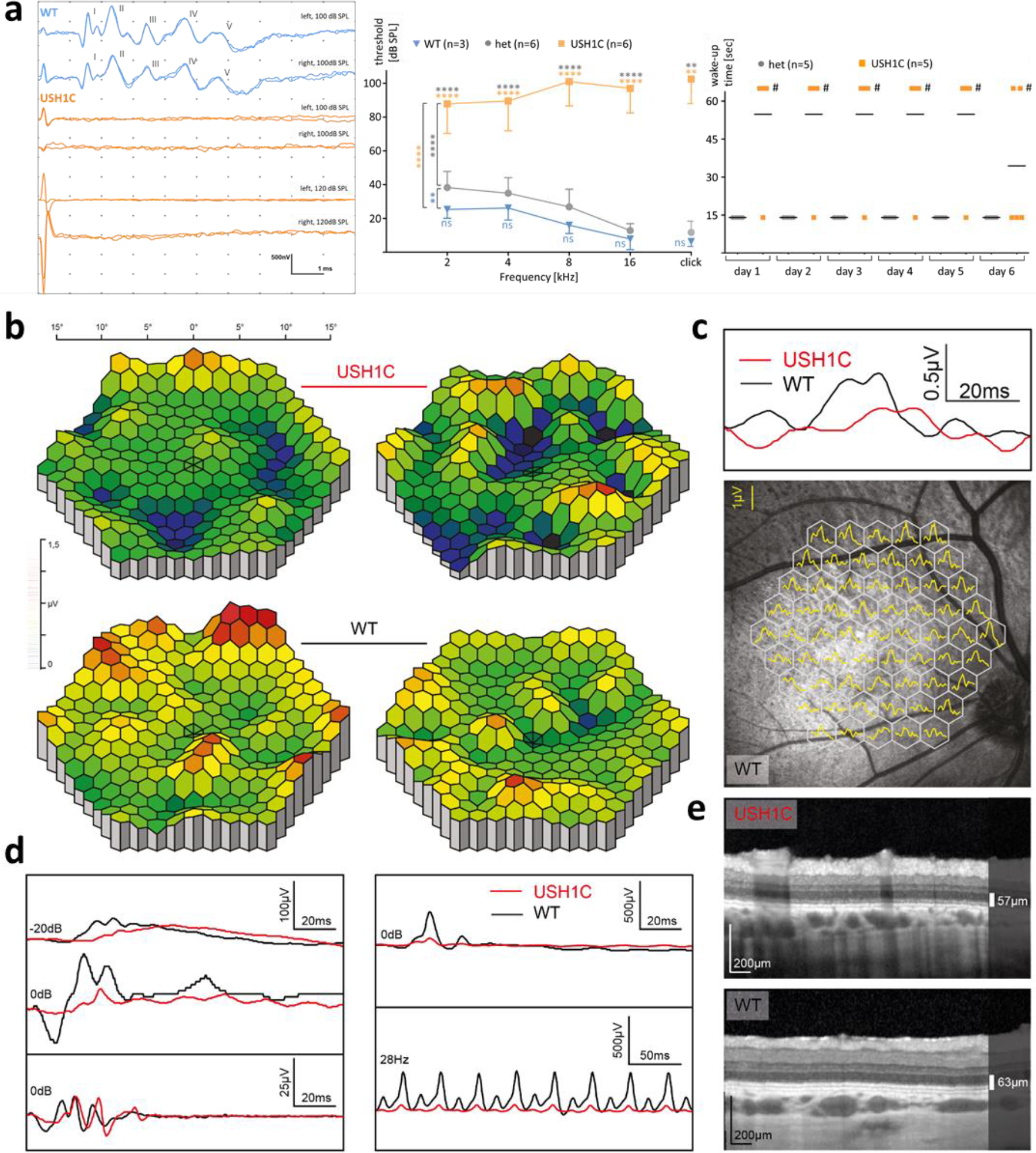
Hearing deficits and impaired vision in USH1C pigs. **(a)** ABR to supramaximal click stimuli revealed profound response at the characteristic peaks I-V in 3-week old WT pigs, but not in USH1C pigs (left). Hearing deficits were more pronounced at higher frequencies in 8-week old USH1C pigs (middle). Differences between WT and heterozygous USH1C^+/−^ (het) were not significant (t-test), for the respective frequencies, but a 2-sided ANOVA test indicated a <0.01 significance (*) overall. P-values for differences between USH1C pigs to WT or heterozygous pigs were <0.0001 (****) at all tested frequencies and <0.01 (**) for a click test. Behavior tests supported impaired hearing in USH1C pigs. In their first days of life, reaction times of sleeping piglets upon calling by the mother sow were measured (right). Piglets waking up immediately entered the box within 15 seconds (Video S7), if not waking up within 60 seconds, piglets were stimulated by nudging (#). Mean values are indicated. In the eye, a retinal dysfunction is evident in multifocal **(b-c)** and Ganzfeld ERG recordings **(d)** with a cone and rod dysfunction that corresponds to a reduced photoreceptor layer in OCT **(e).**

Clinical examination of the eyes in USH1C pigs at 3 and 12 months of age revealed changes comparable to early stages of Usher syndrome. A reduction of cone photoreceptor function was evident in the central visual streak area analysed by multifocal electro-retinography (ERG, Figure 3b-c). Full field ERG demonstrated reduced cone-driven responses at an age of 3 weeks (rod responses were not tested at that age, data not shown). Both rod and cone responses were markedly reduced in USH1C pigs compared to age matched wild type animals at 12 months of age (Figure 3d): USH1C animals showed > 70% reduction in the scotopic standard flash a-wave amplitude ([mean±sd] 21±18 μV in USH1C vs. 79±2 μV in WT animals) and a > 50% reduction in scotopic b-wave amplitude (75±32 μV in USH1C vs. 154±39 μV in WT animals). Responses after photopic standard flash stimuli showed a 50% reduction of a-wave amplitude ([mean±sd] 10±0 μV in USH1C vs. 20±5 μV in WT animals) and a 60% reduction in the respective b-wave amplitude (86±49 μV in USH1C vs. 215±50 μV in WT animals). Confocal scanning laser ophthalmoscopy of the retina and choroid showed no differences between WT and USH1C pigs (data not shown). Interestingly, spectral domain optical coherence tomography (Figure 3e) revealed a reduction in the total retinal thickness in the area of the visual streak, primarily driven by a loss of the outer retinal layers containing the photoreceptor cells (PRC).

### Lack of harmonin in sub-cellular structures disrupts photosensitive disc architecture

In line with data from previous studies (*21, 36*), expression of harmonin was detected in the PRC layer, the outer limiting membrane and outer plexiform layer of WT pigs (Figure S6). More specifically, protein was found in the outer segment (OS) and at its base, as well as in CP of PRC, in cone synaptic pedicles and Müller glia cells (Figure S6b-f). In USH1C pigs, RT-PCR proved correct splicing of the mutated human exon 2 into *USH1C* transcripts in different organs (Figure 4a). Harmonin was consistently lacking at molecular level (Figure 4b) and in tissue sections (Figure 4c), confirming the sufficient premature translational stop by the R31X nonsense mutation. No changes were observed in the expression levels in the retina for other USH proteins, namely SANS (encoded by *USH1G*), whirlin (encoded by *WHRN*, *USH2D*) and myosin VIIa (encoded by *MYO7A*, *USH1B*) (Figure 4d). Up-regulation of the glial fibrillary acidic protein (GFAP) expression was observed in the USH1C retina already at an age of 3 weeks followed by a partial decline (Figure 4e-g). This suggests gliosis, a phenomenon often seen in inherited retinal diseases (*37*) or injury-mediated degradation processes (*38*).

**Figure 4.**
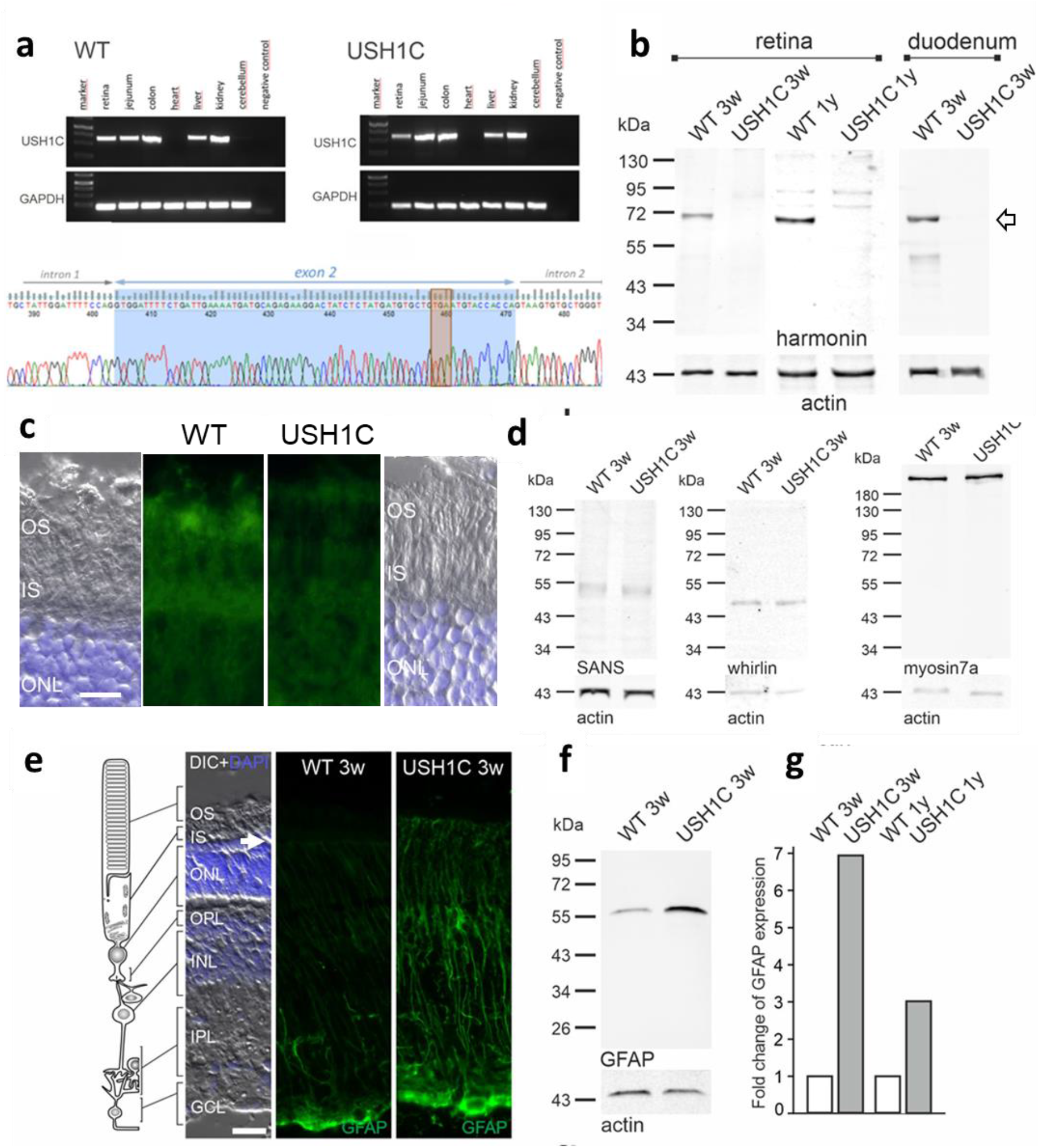
USH1^R31X^ diminish harmonin expression and induce upregulation of GFAP. **(a)** RT-PCR confirms transcription of the *USH1C* gene in WT and USH1C pigs in various organs (upper panel). Sanger sequencing confirms correct splicing of the humanized exon 2 (blue), including the TGA stop-codon (orange box) into the transcripts (lower panel). **(b)** Western blot analysis proved lack of harmonin protein expression in the retina and the duodenum of USH1C pigs at the age of 3 weeks (3w) and 1 year (1y). Arrow indicates expected size of harmonin (isoform a). **(c)** Immunofluorescence staining of harmonin (green) of longitudinal cryosectiorn through the parts of WT and USH1C pig retinas at three weeks of age demonstrated the absence of harmonin staining in the outer segment (OS) of photoreceptor cells. IS, inner segment; blue DAPI staining of nuclear DNA in the outer nuclear layer (ONL). **(d)** Protein abundance of the USH proteins SANS (USH1G), whirlin (USH2D) or myosin7a (USH1B) in the retina seem unaffected at an age of three weeks. **(f)** Immunofluorescence staining of GFAP in Müller glia cells of the WT and USH1C pigs at three weeks of age (3w) which extend throughout almost the entire retina from the outer limiting membrane (arrow) to the ganglion cell layer (GCL) of the retina. The consistent increase of GFAP in expression in the Müller glia of USH1C pigs indicates Müller cell activation and gliosis. OPL, outer plexiform layer; INL, inner nuclear layer; IPL, inner plexiform layer. **(e)** Quantification of GFAP vs actin in Western blots reveals strong 7-fold increase in three week old pigs (3 w) and ~ 3 fold increase of GFAP expression in 1 year (1y) 2 months old pigs when compared to age-matched WT controls **(g)**. Scale bars: C, 10 mm; e, 20 μm.

While the structure and arrangement of PRCs appeared grossly intact in USH1C pigs (Figures 4e, S7a), subcellular analysis revealed specific disruptions in the architecture of compartments comprising harmonin. In line with its proposed scaffold function in synapses (*39*), the loss of harmonin results in significantly reduced diameters of the cone synaptic pedicles, as indicated by measurement of peanut agglutinin (PNA) pattern (Figure S7b) as well as by the direct synapse width in transmission-electron microscopy (TEM) (Figure S7c). We also found significant increases in length of the connecting cilium (CC) in rod PRCs, as indicated by fluorescence microscopy of the ciliary marker GT335 (Figure 5a-c) as well as by TEM examination of ultrathin sections (Figure 5d, f). Consistent with this, lacking of harmonin led to the elongation of primary cilia in primary dermal fibroblasts derived from USH1C pigs and a USH1C^R31X/R80fs^ patient after induction of ciliogenesis (Figure 5f-h). Most striking, however, was the deprivation of rod OS architecture in PRC. While this is normally characterized by a strictly horizontal stacking of parallel photosensitive membrane discs (Figure 5i) (*40*), we detected vertically orientated membrane discs (Figure 5j) as early as at three weeks of age. This aberrant structuring became even more prominent at 12 months when the stacking of discs was further infringed by interstitial gaps (Figure 5k-m). Additionally, vesicle-like structures were observed at the OS base (Figure 5l). Interestingly, and in contrast to rods, OS architecture in cones remained normal throughout the first year of life (Figure S7a).

**Figure. 5.**
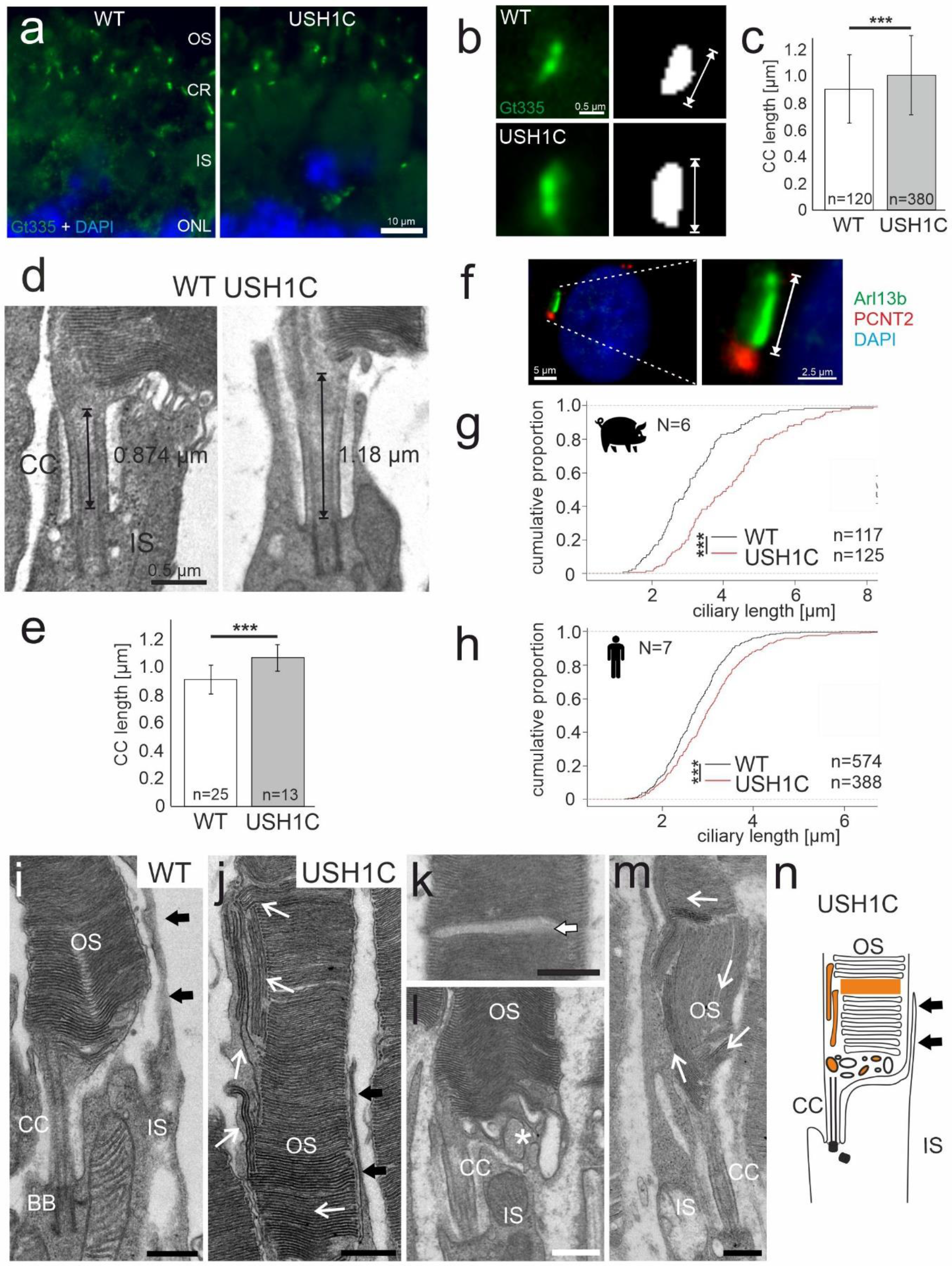
Changes in ciliary structures affect PRC outer segment architecture. **(a)** Immunostaining of the ciliary marker GT335 (green) in cryosections through ciliary regions of PRCs of WT and USH1C retinas. Length measurements of stained cilia at high magnification **(b,c)** and in longitudinal sections through the ciliary regions of rod PRCs by TEM **(d,e)** revealed significant longer connecting cilium (CC) in USH1C PRCs in both cases **(c, e).** Numbers in box plots indicate number of measured cilia. DAPI staining (blue). In primary dermal fibroblasts derived from USH1C pigs **(f,g)** and from a human USH1C^R31*/R80fs^ patient **(h)**, primary cilia were stained for ARL13B (ciliary shaft, green) and PCNT2 (ciliary base, red). Consistently, mutated *USH1C* caused longer primary cilia when 117 WT vs 125 USH1C porcine cells **(m)** or 574 WT vs 388 USH1C human cells **(n)** were compared. Two-tailed Student’s t tests, \*\*\**p*≤0.001. **(i-m)** TEM of longitudinal sections through rod PRCs of WT and USH1C pigs. At three weeks of age the overall architecture, appears similar between WT (i) and USH1C **(j,l)** including the abundance of calyceal processes (black arrows). In USH1C rods, however, vertically oriented membrane discs (**j**, white arrows) appeared in the outer segment (OS) and vesicles are found at the OS base (**l**, asterisk). At 12 months of age, in USH1C OSs distorted discs are found (**m**, white arrows) and interstitial gaps appeared in disc stacks **(k)**. **(n)** Schema of a USH1C rod summarizing OS architecture anomalies highlighted in orange. Scale bars: i, 0.6 μm; j, 0.75 μm; k, 0.55 μm; l, 0.5 μm; m, 0.85 μm.

### USH1C pigs facilitate testing of therapeutic approaches *in vivo* and *ex vivo*

As adeno-associated viruses (AAV) comprise the most popular vector to deliver nucleic-acid-based therapies into a patient, we tested the principal applicability of AAV *in vivo*. AAV8, AAV9 and Anc80 expressing eGFP under control of a CMV promoter (*41, 42*) were injected into the sub-retinal space of the eye in WT pigs (Figure 6a). While AAV8 transduced only photoreceptor cells efficiently, AAV9 and Anc80 transduced also Müller glia cells, indicating that the target specificity in the eye can be controlled by the appropriate choice of the AAV capsid. No relevant other off-target expression of eGFP was noted for the tested AAVs.

**Figure 6.**
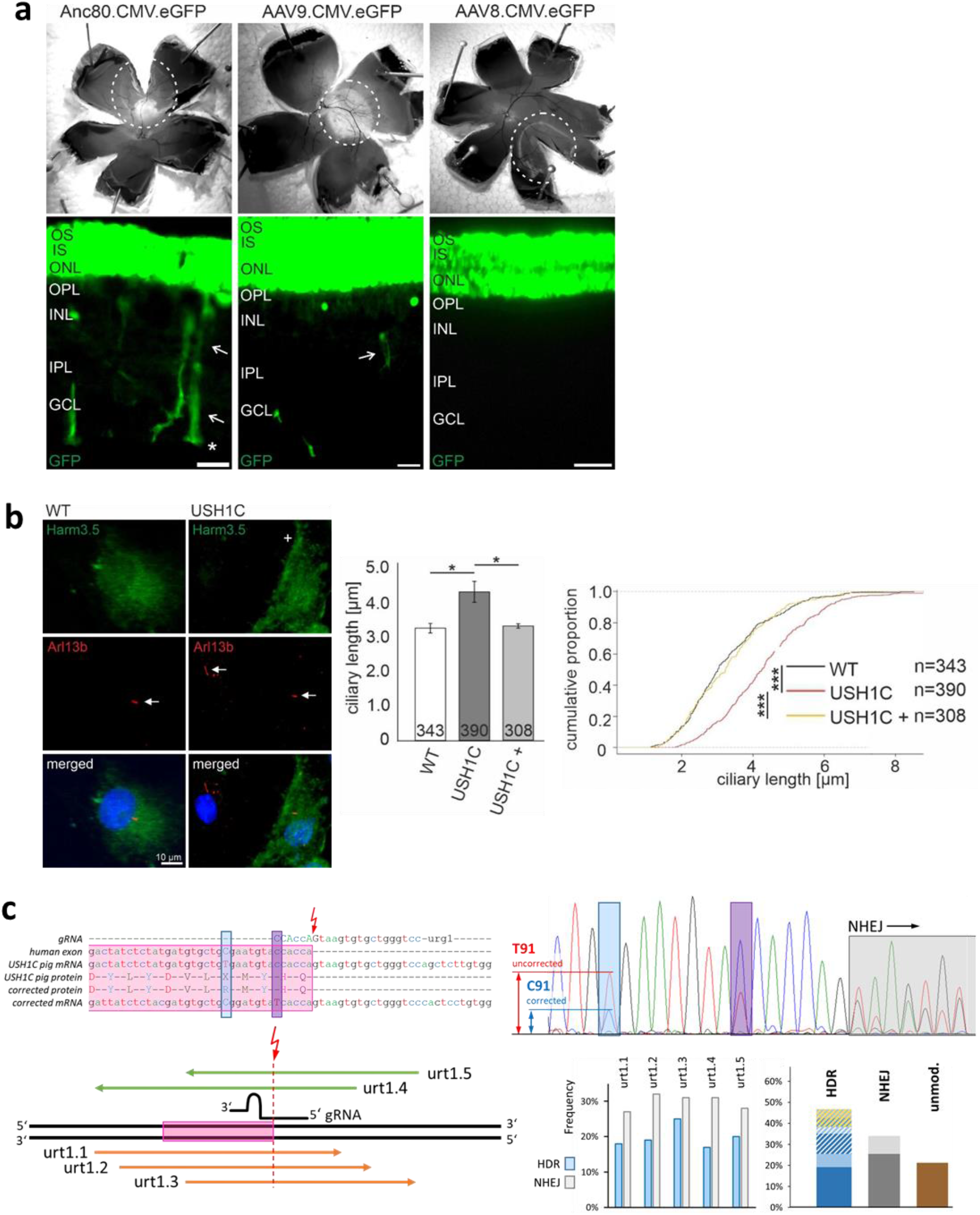
Therapeutic approaches. **(a)** Expression analysis of AAVs. AAV8, AAV9 and Anc80 expressing eGFP were injected subretinally into the eyes of WT pigs *in vivo*. Epifluorescence of dissected eyes demonstrates eGFP fluorescence in the bleb region of whole mount preparations of all three AAV samples (upper row). Epifluorescence of longitudinal cross-sections revealed abundant eGFP expression in the photoreceptor cell layer (OS, IS and ONL) throughout the treated bleb region for all AAV and in Müller glia cells upon transduction with AAV9 and Anc80 (arrows) (lower row). Scale bares, 25 μm. **(b)** Rescue of the ciliary length phenotype in dermal primary fibroblasts of USH1C pigs by human harmonin_a1 expression. Immunofluorescence analysis of USH1C cells does not show harmonin expression in untransfected cells (right panel, left cell, +) and abundant expression of harmonin in transfected cells (right panel, right cell). Quantitative analysis of primary ciliary length reveals significant decrease of ciliary length in USH1C+ fibroblasts towards the ciliary length in WT. Harm3.5, harmonin staining; Arl13b, primary ciliary shaft staining (arrows). Two-tailed Student’s t test, *p≤0.05, **p≤0.01. **(c)** In CRISPR/Cas-mediated gene repair, the cutting site (red arrow) of the most favorable gRNA was directly at the transition from exon 2 (pink) to the downstream intron (upper left). Distinct ssODN binding to the sense (green) or anti-sense strand (orange) were designed to correct the non-sense mutation to an Arg codon (blue) and to introduce a blocking mutation at the PAM site (magenta) (lower left). After determining the frequency of HDR and NHEJ in mixed cell populations (right panel, Figure S6), single cell clones were examined for the allelic occurrence of correcting mutations alone or in company with NHEJ (n=101) after treatment with urt1.3 (lower right). The frequency of HDR was discriminated for homozygous (dark blue) or heterozygous repair (light blue). Cell clones in which HDR was accompanied with NHEJ are shaded in blue/white if the mutation appeared in the intron and blue/gold if mutations are presumably disruptive for the ORF. Cell clones with sole NHEJ events are shown in dark or light grey, when they appeared on both or only one allele. The frequency of unmodified clones is depicted in brown.

For exploring the potential of harmonin restoration by gene therapy we transfected a vector encoding the human *USH1C* splice variant a1 (Nagel-Wolfrum et al., unpublished data) into primary skin fibroblasts and determined the length of the primary cilium (Figure 5f-g) as therapeutic parameter (Figure 6b). After restoration of harmonin abundance in USH1C cells, the mean ciliary length decreased to the level of WT cells.

The capability for correcting the causative *USH1C^C91T^* mutation in the humanized exon 2 by gene repair was evaluated by applying distinct constellations of human-specific gRNA and repair oligo-nucleotides into PKCs (Figures 6c, S8). The efficacy of transforming the nonsense codon TGA into an Arginin-encoding CGA by homology directed repair (HDR) reached 40%. Clonal selection revealed that 19% of the cells showed HDR on one allele and 22% on both alleles. NHEJ was seen in 32% and 8%, respectively. Occasionally, correction of the R31X codon was accompanied by additional mutations, either small insertions/deletions or single nucleotide exchanges. For the specific positioning of the gRNA, most of these modifications appeared within intron 2 and, therefore, would not affect the correct transcription and translation of *USH1C*.

## Discussion

Demonstrating the combination of hearing deficits, vestibular dysfunction and vision loss, the USH1C pig is the first animal model to reflect the full syndromic phenotype in human USH1 patients. The early onset of symptoms makes it suitable for studying the basic mechanisms of the disease as well as for evaluating strategies of therapeutic intervention. The persistence of a largely intact retinal architecture throughout the first year of life provides a wide therapeutic window and the potential to revert the pathogenesis by targeting dysfunctional, but still viable photoreceptor cells. The specific design of the genetic modification allows the examination of a broad diversity of therapies, including classical additive gene therapy, human-specific gene repair, translational read-through or cell replacements. After establishing breeding populations and demonstrating therapeutic efficacy and successful application of AAV vectors (Figure 6) the USH1C pig can now be used for pre-clinical examination *in vivo*, similar to recent ventures in genetically engineered pig models (*43–45*). In such studies, housing USH1C pigs under standardized conditions will minimize environmental confounding, whereas the pronounced genetic diversity of conventionally produced pigs needs consideration. While presenting a statistical challenge for pre-clinical proof of principle studies, this heterogeneity is, of course, much more likely to reflect the scenario in clinical trials and might thus lead to a more robust predictive power of the model in the translation of new therapeutic strategies (*46*).

The manifestation of an ocular phenotype in the USH1C pig is striking, taking into account the multiple efforts that have been made in creating such models in other species with so limited success (*6*). Our results reinforce some of the multiple sentiments that have been postulated as reasons for the lack of a vision phenotype in rodents. The onset of retinal dysfunction in pigs within the first year of life argues against the notion that the limited life span of mice prevents developing a similar phenotype in the mouse eye. While a potentially distinct function of the USH proteins in the retinae of different species cannot be readily excluded, the degree of sequence conservation of harmonin and its immediate interaction partners is remarkable (Figures 1, S9), insinuating a conserved and consistent nature of the USH1 interactome across species.

Crucially, porcine harmonin appears localized to CP (Figure S6d) which are thought to stabilize the OS structure in PRC of diverse species, but not in rodents (*5*). A recent publication demonstrated the absence of CP in cone PRCs and a pronounced outgrowth of disc membranes at the base of rod OS upon partial knock-down of the USH1F protein pcdh15 in the claw frog (*10*). While the complete lack of harmonin in pig did neither show outgrowth of disc membranes nor affect the formation of CP to a similar extent, it explicitly disturbed the photosensitive disc architecture in rods (Figure 5i-n). The changes in OS architecture, disc arrangement and CC dimensions (Figure 5) upon abolishing harmonin function as an actin-binding protein ^21^ confirms the driving nature of the actin cytoskeleton in disc neogenesis and suggest relevance of harmonin in disc stacking (*47, 48*). Furthermore, we have identified elongation as a common phenotype of CC in PRC of USH1C pigs and primary cilia in dermal fibroblast of both, USH1C pigs and human patients (Figures 5a-h, 6b). This is in line not only with the close association of the ciliary actin system with axonemal microtubules in PRCs (*47, 49*), but also with the functional role of the actin system in maintaining the organization of the ciliary axoneme in general (*50*). The elongation of cilia in harmonin-deficient cells is probably due to the dysregulation of the ciliary actin cytoskeleton (*51*) which has been also observed in other neuronal degenerative diseases (*52*). Although precise mechanistic insights into the role of harmonin in cilia function require further studies, we provide clear evidence that USH is a true retinal ciliopathy in its fundamental aspects (*53*).

Remarkably, the significantly disturbed photosensitive disc architecture in rod OSs and the ciliary aberrations (Figure 5) are associated with a significant (c. 70%) reduction in rod-derived ERG responses (Figure 3d). The less pronounced decline in cone-derived ERG (c. 50%) and the reduced width of cone synaptic pedicles (Figure S7b, c) indicate the early onset of cone-specific alterations. Both, structural and clinical observations thus support the notion of a rod-cone dystrophy phenotype in USH1C pigs, reflecting the situation in human patients quite well (*35, 54*). The early onset of PRC malfunction was in line with the up-regulation of GFAP in Müller glia cells (Figure 4e-g) and with reports of visual deficits in early childhood of USH patients (*55*). Interestingly, the changes in subcellular morphology, the reduction in electrophysiological PRC function and the observed changes in visually guided behaviour come along with only very mild effects on overall retinal morphology. It needs to be considered, however, that the retinal phenotype in USH1 patients varies widely, ranging from intact central retinal architecture and excellent visual acuity at 45 years to advanced central retinal damage and legal blindness by their early 30ies (*56*). Such a phenotypic heterogeneity is observed in most if not all forms of inherited retinal diseases and is thought to arise from genetic or environmental confounding factors or a combination thereof (*57*). The possibility to examine the USH1C pig model under standardized conditions will help to discern the effect of the mutation from other influencing factors.

For the different ageing processes in pig and human, a direct correlation of ages between the species is misleading. Instead, taking into account the manifestation of fertility at 5-8 months in pigs, compared to 12-15 years in human beings, a 12-months old pig may fairly correspond to a biological age of 15-25 years in humans (http://www.age-converter.com/pig-age-calculator.html). Accordingly, the phenotype in the USH1C pig appears to reflect patients well within the published spectrum of disease severity and progression. As consequence, translating clinically established end-points such as ERG or OCT from human to pig will support the assessment of future pre-clinical studies in our model. Vice versa, the multi-disciplinary examination of the model described here defined novel disease hallmarks, which may give rise to alternative endpoints for future clinical trials. The value of additional read-out parameters is corroborated by a pivotal study documenting the success of *voretigene neparvovec* on basis of motion analysis in the multi luminance mobility test (MLMT) as primary endpoint (*58*).

The specific wave-length-sensitivity of cones (*59*) and the complex interaction of vision with other sensory organs required substantial effort in the investigation of visually guided behaviour. Parcour tests were conducted under distinct light conditions, but calculated luminance levels in the pig eye suggested experiments took place at the lower and upper limit of mesopic sensing (*60*). Albeit this prevents a clear discrimination of rod- and cone-based vision in the motion analysis, essential conclusions can be drawn. First, the similarity of core parameters in the hurdle parcour between USH1C animals and WT controls in the dark confirms that pigs substantially rely on non-vision sensing and denies an exclusive or dominant effect of vestibular dysfunction on movement. Second, the clear differences in the light and the characteristic difficulties that USH1C pigs have in the recognition of defined objects (Figures 2a-c) document their substantial problems in 3D orientation. The faster and smoother trajectory (Figure 2d) of WT control pigs further demonstrates how a normal visual function promotes effective clearance of obstacles and „elegant“ motion.

Taken together, the generation of an USH1C pig model has substantially improved our molecular understanding of the disease. We now have several lines of evidence how harmonin supports PRC function and the particular integrity of the OS. The early onset of basic defects in the retina and the impairment during the first year provides an ideal foundation for following the natural history of USH1C. Combining quantitative motion analysis with clinical examination tools will promote the definition of read-out parameters in pre-clinical studies and in clinical trials for treating Usher syndrome, but also patients with other progressive blinding diseases.

## Material and Methods

### Regulatory statement

Work on USH pigs has been conducted under the supervision of the responsible regulatory authorities: the Regierung von Oberbayern has approved animal experiments involving somatic cell nuclear transfer, maintenance of pigs and longitudinal monitoring at LMU Munich under the file number AZ 02-17-136. The State Veterinary Administration of the Czech Republic approved animal experiments on maintenance, sub-retinal intervention and longitudinal monitoring at IAPG Libechov under the experimental protocol number 75/2019.

### Methods

Transgenic modification of pigs and their generation has been conducted as described in (*23, 24*); postnatal care has been described (*27*). Sampling as well as morphological and molecular analysis was done according (*61*). Intervention and monitoring has been described previously (*62*). For more details and description of behaviour tests see the supplementary material section.

## Author contributions

**S.G.** contributed to molecular analysis and breeding of animals, designed and performed behaviour tests and contributed to writing of the manuscript. **J.S.** contributed to molecular and morphological analysis, and ciliation analysis in primary fibroblasts. **K.A.W.** contributed to tissue sampling and processing, imaging (IF and SEM), data analysis and interpretation. **Z.E.** contributed to breeding attempts sub-retinal intervention and ABR measurements. **H.A.** evaluated Gene Repair. **A.Bä.** managed intervention on pigs and anaesthesia. **P.R.-V.** contributed to Gene Repair study. **J.Pl.** contributed to genetic modification. **V.A.** contributed to quantitative analysis of the synaptic phenotype in cones. **T.A.** managed intervention on pigs. **M.V.** performed subretinal injection. **G.S.** performed trajectory analysis. **G.D.** contributed to animal management and tissue sampling. **W.H.** performed data analysis. **B.K.** contributed to Somatic Cell Nuclear Transfer and performed Embryo Transfer. **M.K.** & **V.Z.** performed Somatic Cell Nuclear Transfer. **J.L.** performed ciliation analysis in primary fibroblasts. **A.F., S.S.** & **J.Po.** performed ABR measurement. **A.Bl.** conducted pathological examinations and tissue preparation. **A.D.** contributed to ERG measurement. **H.M.-S.** assisted with tissue dissection and cochlear preparation. **K-L.L.** & **E.W.** contributed to the design of the study and the writing of the manuscript. **L.H.V.** provided AAV vectors. **K.N.W.** contributed to the design of the study, interpretation of data and writing of the manuscript. **J.M.** contributed to the design of the study, orchestrated breeding attempts, sub-retinal intervention and ABR measurements. **M.D.F.** led the ophthalmological characterization of animals, performed the vitreoretinal surgery and wrote the manuscript. **U.W.** designed and coordinated the study, performed molecular and morphological analysis, interpreted data and wrote the manuscript. **N.K.** designed and coordinated the study, performed bio-informatic and molecular analysis, interpreted data and wrote the manuscript.

## Acknowledgements

This work has been supported by the FAUN foundation, USHER2020, and Foundation Fighting Blindness (to: E.W. N.K. & U.W. (TA-GT-0316-0693-JGU); K.N.W. & U.W. (TA-GT-0316-0694-JGU) and to U.W. (PPA-0717-0719-RAD) as well as by the German Research Council within SPP2127 “Gene and cell-based therapies for neuroretinal degeneration” (to: M.D.F. (FI 2336/1-1); H.M.-S. (MA 6139/3-1); K.N.W. (KA1398/1); U.W. (WO248/9); N.K. (KL 2405/2-1). We are grateful to Elisabeth Sehn, Ulrike Maas, Gabi Stern-Schneider and Christian Eckardt for their excellent technical support, to Martin Pavlik and Alena Havrdova for excellent animal caretaking, to Marold Handl for necropsy assistance. Many thanks also to the Institute of Pathology, LMU Munich for providing facilities and infrastructure for necropsies.

## Competing interests

The authors declare to have no conflict of interest.

## Supplementary Figures

**Figure S1.**
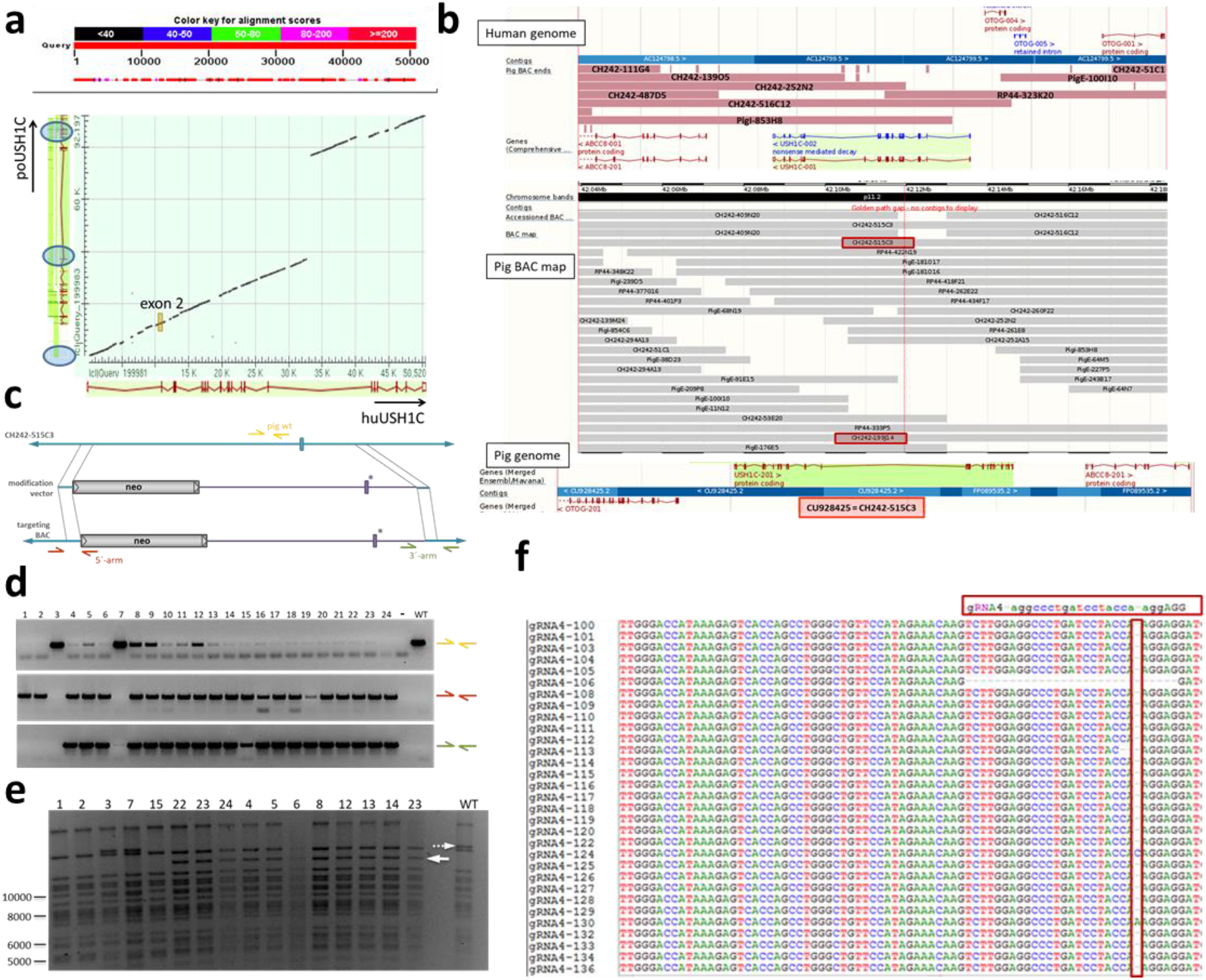
Modification strategy. **(a)** The annotation of the porcine *USH1C* gene was grossly confirmed by BLAST search with the b3-splice variant of human *USH1C* mRNA (GenBank no. NM_153676.4), including exon 2. Exons not correctly defined in the pig genome are marked in blue circles. The position of exon 2, containing the R31 codon, is marked in the coverage plot by a yellow box. **(b)** Based on the matches of pig BAC end-sequences to the human genome and the pig reference genome Sscrofa 10.2, the *USH1C* locus was defined in the Pig-Pre BAC map and BAC covering the entire *USH1C* locus were purchased (red boxes). Clone CH242-515C3 was confirmed by BAC end sequencing to represent the GenBank no. CU928425. **(c)** A modification plasmid comprising the intended 1.5kb human *USH^R31X^* segment (magenta) and a floxed neo selection cassette was introduced into BAC CH242-515C3 by bacterial recombineering. Screening for correct recombination was performed with end-point PCR for the 5′-end (brown), the 3′-end (green) and the loss of the original porcine sequence (yellow). **(d)** Bacterial clones comprising the desired constellation were further confirmed by BAC restriction digest with XbaI **(e)** or SpeI (not shown). Introduction of the modification changes the digest pattern only slightly. In the XbaI digest, an 18779bp band in the WT BAC (dotted arrow) is modified to a 14194bp band (arrow) and a 5020bp band (not shown). **(f)** gRNA were tested in pig primary cells for their efficacy to introduce mutations via NHEJ by Sanger sequencing of cloned PCR products.

**Figure S2.**
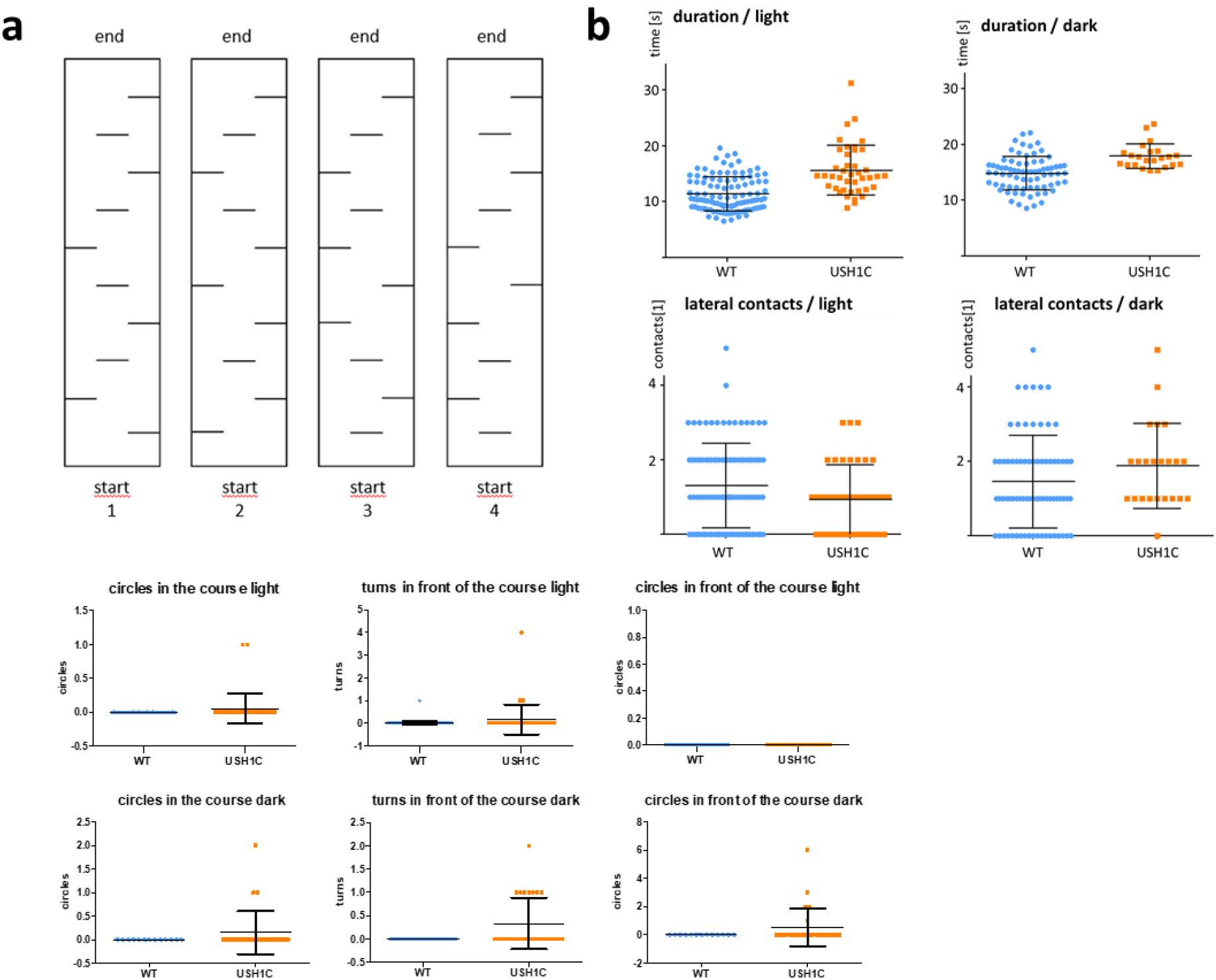
Barrier course. **(a)** Shields were placed in consistent distance, but in a different order, either on the left, the right or in the center of the course. **(b)** Indicative parameters were evaluated for USH1C pigs vs WT controls, either in the dark (average of 2.9 lux) or under normal light conditions (average of 136 lux).

**Figure S3:**
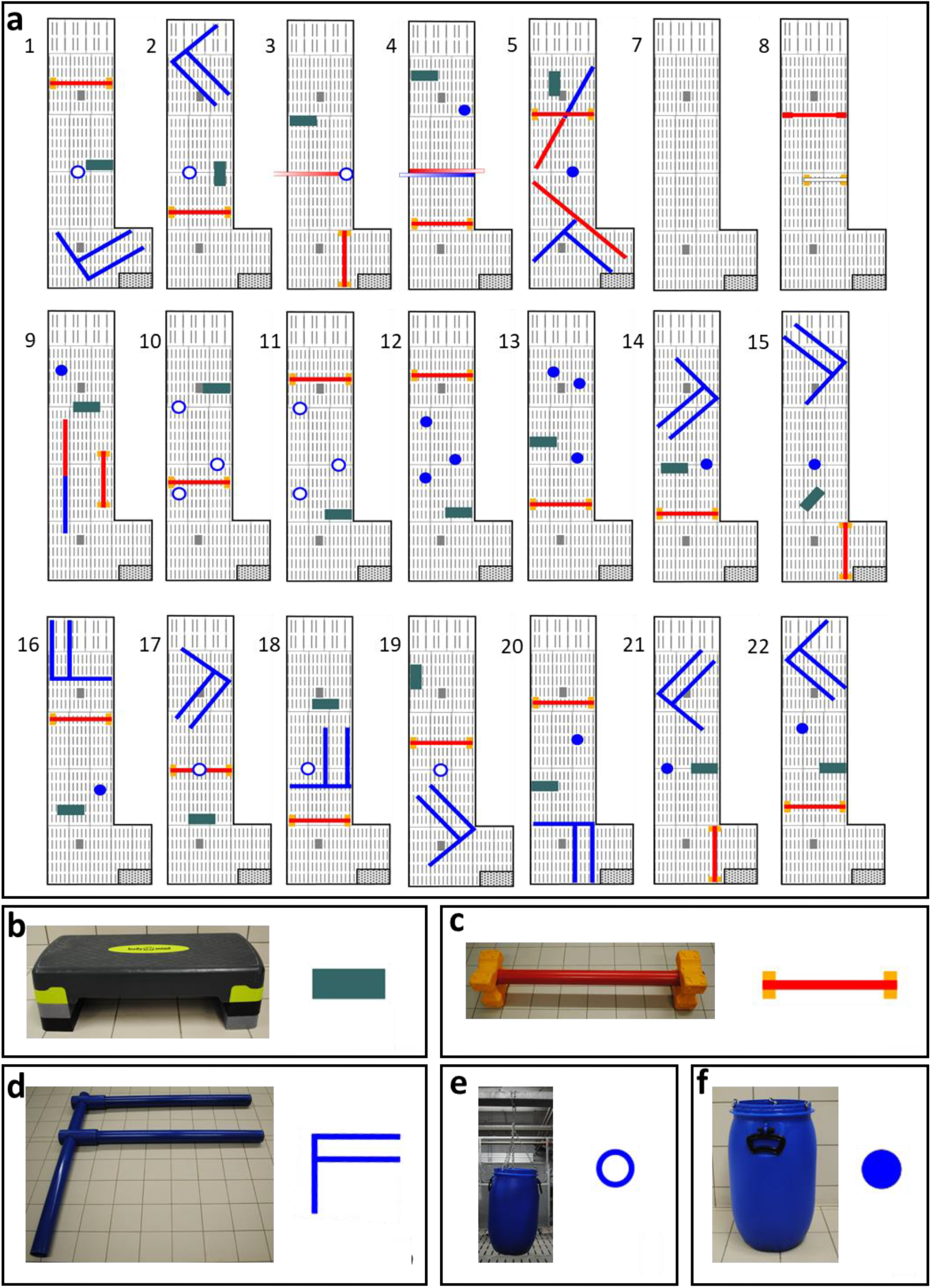
Obstacle course set-up. **(a)** obstacles were used in different constellation and different order at 22 test days. **(b) - (f)** photographs and symbols of most frequently used obstacles: step board (b), cavaletti (c), “F“ formed out of poles (d), barrel hanging from the ceiling (e), barrel placed on the floor (f).

**Figure S4:**
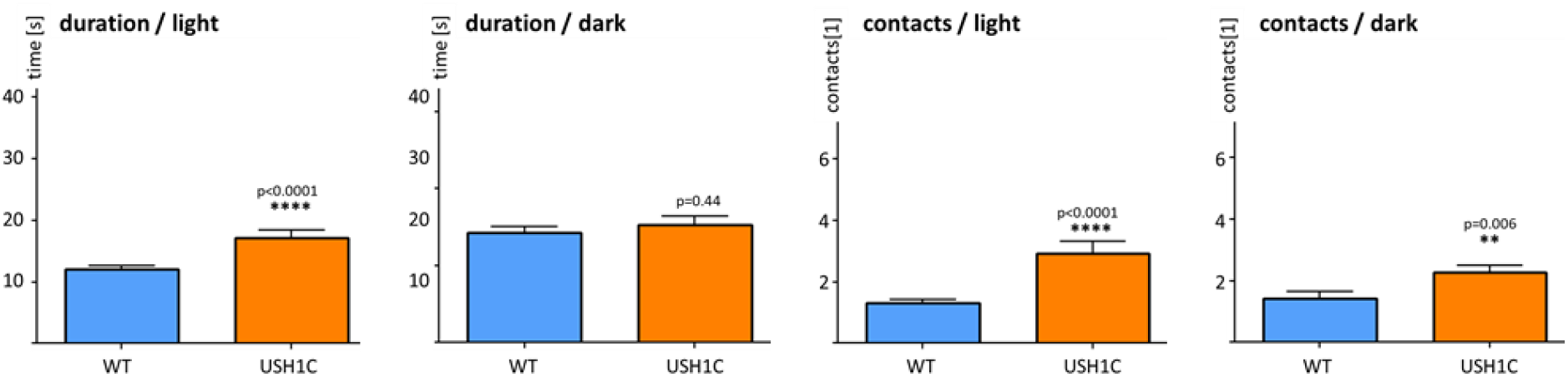
In-depth statistical analysis of obstacle course. Data were checked for consistency and normality. Fisher’s Exact test or Pearson’s test were used to analyze cross tabulations. Generalized linear models with Poission distribution, Median tests, bootstrap-t tests based on 5000 Monte Carlo simulations, t-tests with and without the assumption of homogeneity, Mann-Whitney U tests were used to test continuously distributed variables. All reported tests were two-sided, and p-values < 0.05 were considered statistically significant.

**Figure S5.**
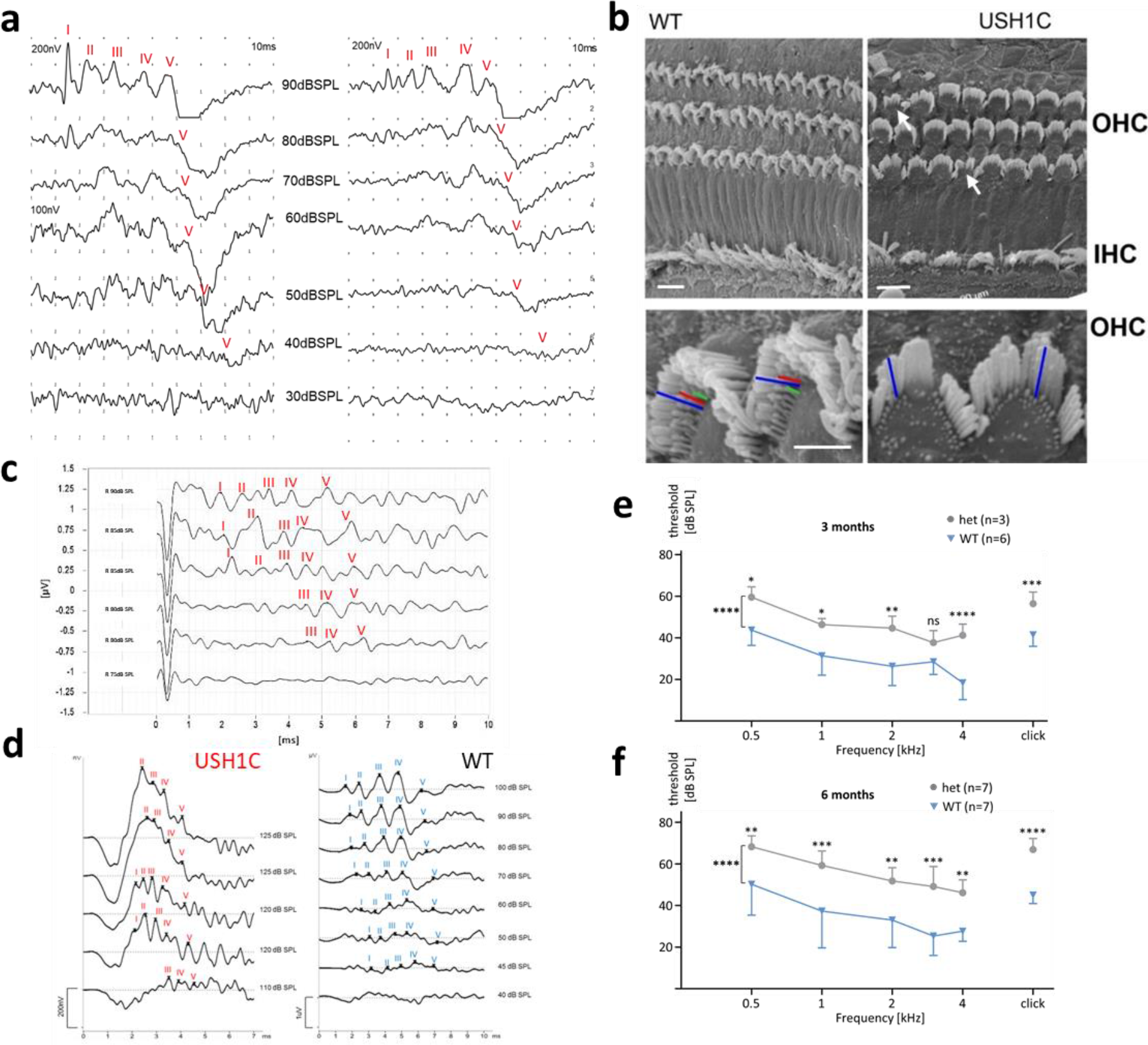
Impaired auditory system in USH1C pigs. **(a)** ABR measurements were conducted on two WT pigs to determine the threshold for sufficient detection of response peak V to click stimuli. **(b)** Scanning electron microscopy of the inner ear in 3-week old WT and USH1C pigs reveals alterations in the hair bundle arrangements of outer (OHC) and inner cochlear hair cells (IHC) (arrows). USH1C OHC lack stereocilia rows in the hair bundles, indicated by colored lines at higher magnification. Scale bars, upper panel 10 μm; lower panel: 5 μm. **(c)** and **(d)** show exemplary response curves for determining sound pressure threshold levels in an 8-week-old and a 2-year-old USH1C animals, respectively. ABR on animals at an age of 3 months **(e)** and 6 months **(f)** indicate an increased threshold of heterozygous USH1C^+/−^ pigs, compared to WT littermate controls. Statistical examination was carried out by t-test for each frequency, a 2-side ANOVA test was performed over all frequencies.

**Figure S6.**
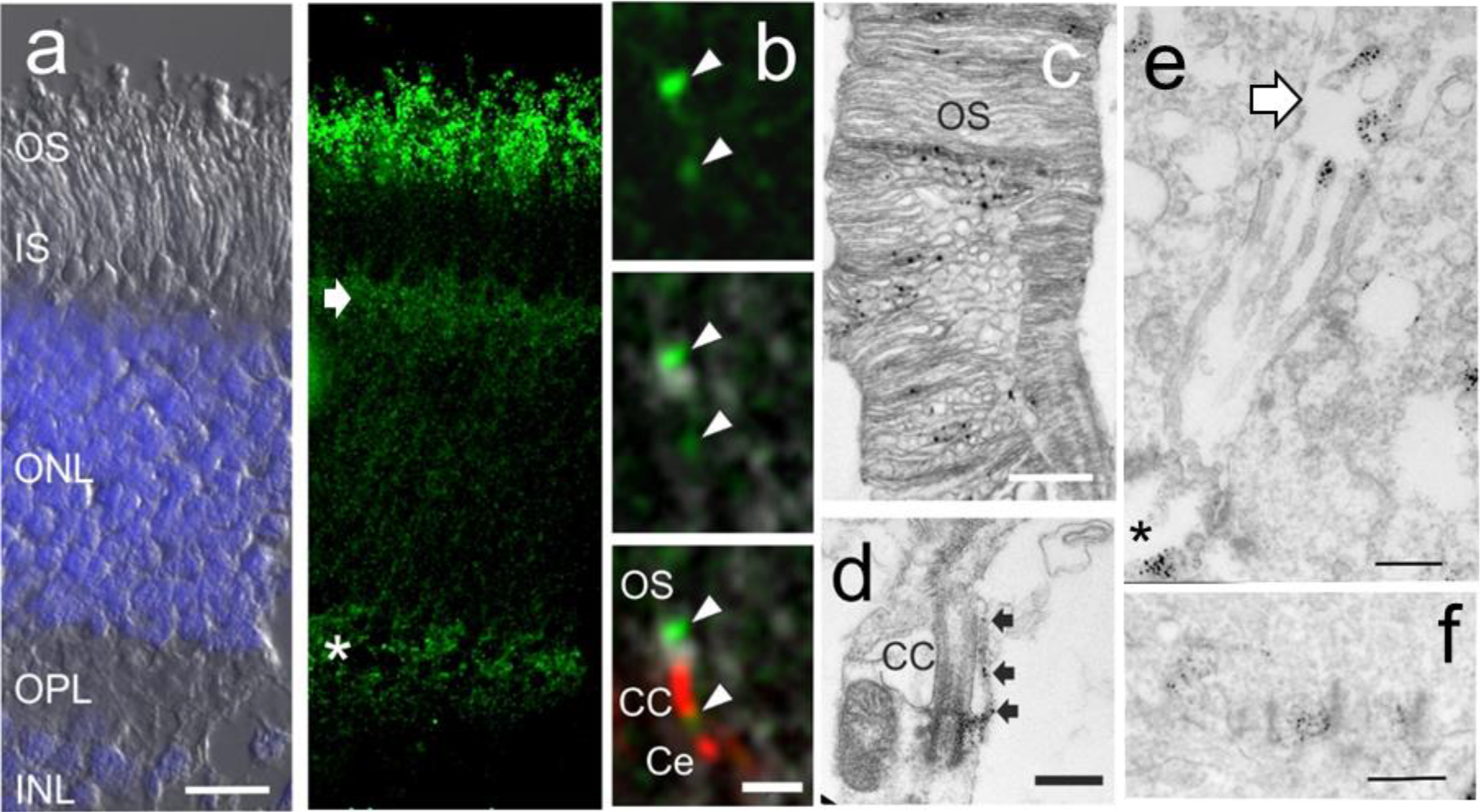
Localization of the USH1C protein harmonin in WT retina. **(a)** Immunohistochistochemical localization of harmonin in longitudinal cryosections through the WT porcine retina. Harmonin (green) is present in the layer of photoreceptor outer segment (OS), the outer limiting membrane (*arrow*) and in the outer plexiform layer (PLA, *asterisk*), where photoreceptor synapses are located. Left image represents DIC image super exposed with DAPI staining of nuclei in outer nuclear layer (ONL) and inner nuclear layer (INL). IS, photoreceptor inner segment; IPL, inner plexiform layer. (**b)** High magnification of immunostained harmonin (green) and centrin (conneting cilia (CC)/centriole (Ce) marker) (red) at base of outer segment (OS) and (*arrowheads*) of WT photoreceptor cells. (**c)** Immunoelectron microscopy of harmonin at the OS disks of a WT rod photoreceptor cell (**d)** Immunoelectron microscopic staining of harmonin in a calyceal process (*arrows*) and the ciliary base of a WT photoreceptor cell. (**e)** Harmonin in immunostaining in Müller glia cells: microvilli tips (*arrows*); at cell adhesion in OLM (*) and (**f)** in cone synaptic pedicles. Scale bars: a, 10 μm; b 1 μm c 0.5 μm d, 0.5 μm e, f: 0.5 μm

**Figure S7:**
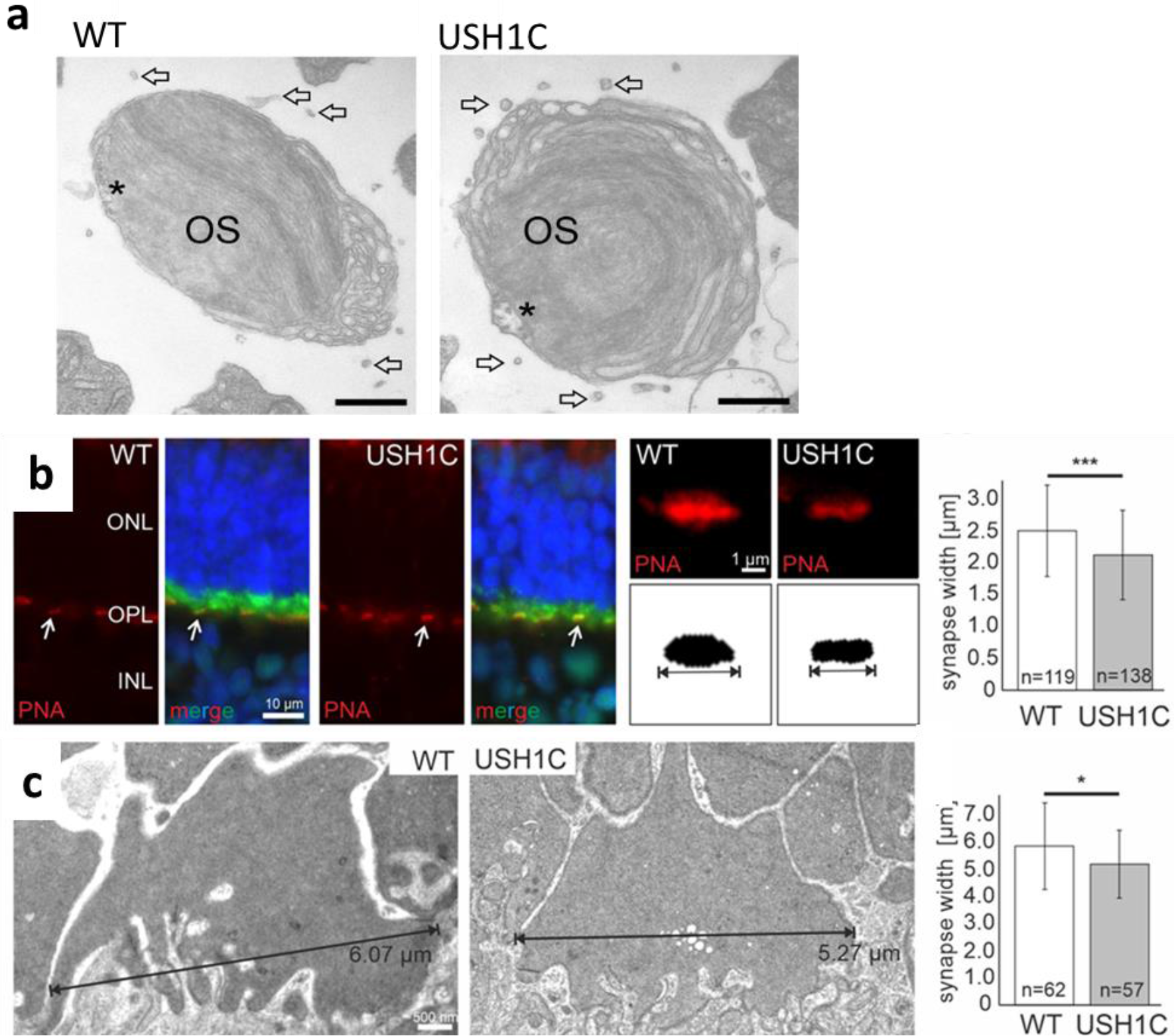
Analysis of outer segments **(a)** and synaptic pedicles of cone photoreceptor cells of WT and USH1C pigs **(b,d)**. **(a)** TEM analysis of horizontal cross sections through the outer segments (OS) of cones reveal persistance of calyceal processes (arrows). Projection of the axoneme into OS is indicated by asterisks. Scale bar: 400 nm. **(b)** Quantitative fluorescent microscopic analysis of cone synapse. **Left panel:** longidudinal sections through WT and USH1C pig retinas stained for the pre-synaptic marker PSD-95 (green) and by fluorescent peanut agglutinin (PNA, red) for cone synaptic pedicles (white arrows) and counter-stained by DAPI for nuclear DNA (blue). ONL, outer nuclear layer; OPL, outer plexiform layer; INL, inner nuclear layer. **Middle panel:** higher magnification of a PNA-stained cone synaptic pedicles. Synapse width was determined as the maximum extension of consistent PNA signals. **Right panel:** Box plot of cone synaptic pedicle width in WT and USH1C pigs revealing significantly reduced synapse width in USH1C cones. **(h)** TEM analysis of the width of cone synaptic pedicles of WT and USH1C pig retinas. Box plot quantification of synaptic width confirmed the significantly reduced width of cone synaptic pedicles in USH1C pigs. Scale bar: 500 nm. Two-tailed Student’s t tests, \**p*≤0.05, \*\*\**p*≤0.001

**Figure S8:**
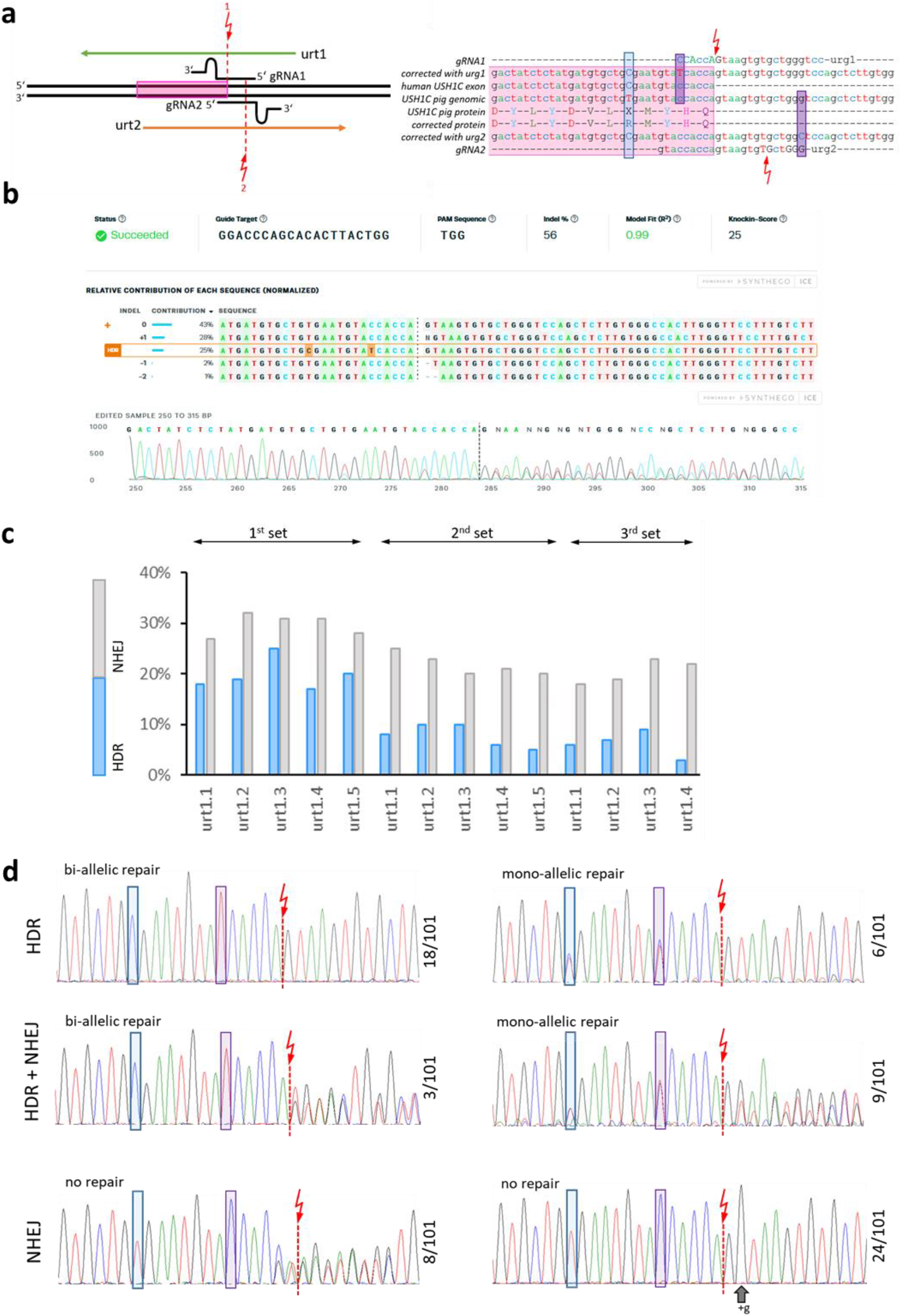
Gene Repair. **(a)** In an initial approach the 2 oppositely oriented gRNA1 and gRNA2 and repair oligonucleotides urt1 and urt2 were tested. Exon 2 is marked by a pink box. The cutting sites of the Cas9 are shown as red arrows and the distinct positions at which the respective oligo-nucleotides should introduce blocking mutations are indicated by magenta boxes. The position of T91 and its corrected variant C91 is indicated by a blue box. **(b)** Upon co-transfection into primary cells from USH1C pigs, PCR products from mixed cell clones were analyzed for NHEJ and HDR with the ICE CRISPR Analysis Tool. **(c)** Optimization was performed with gRNA1 and five distinct repair oligo-nucleotides in three independent experiments. The rate of HDR and NHEJ were determined as in (b). **(d)** Single cells clones were generated from the pool nucleofected with gRNA1 and urt1.3 and analyzed by Sanger-Sequencing of PCR products. A diverse pattern of distinct modifications, comprising HDR, NHEJ or combined variants, at the modified *USH1C* alleles was observed. The cutting site of Cas9 as well as the correcting and blocking mutation are indicated as in (a). The frequency of the respective pattern is given on the right side of the electropherogram, revealing that 36 out of 101 cell clones showed a correcting T91C transmission at least at one allele.

**Figure S9:**
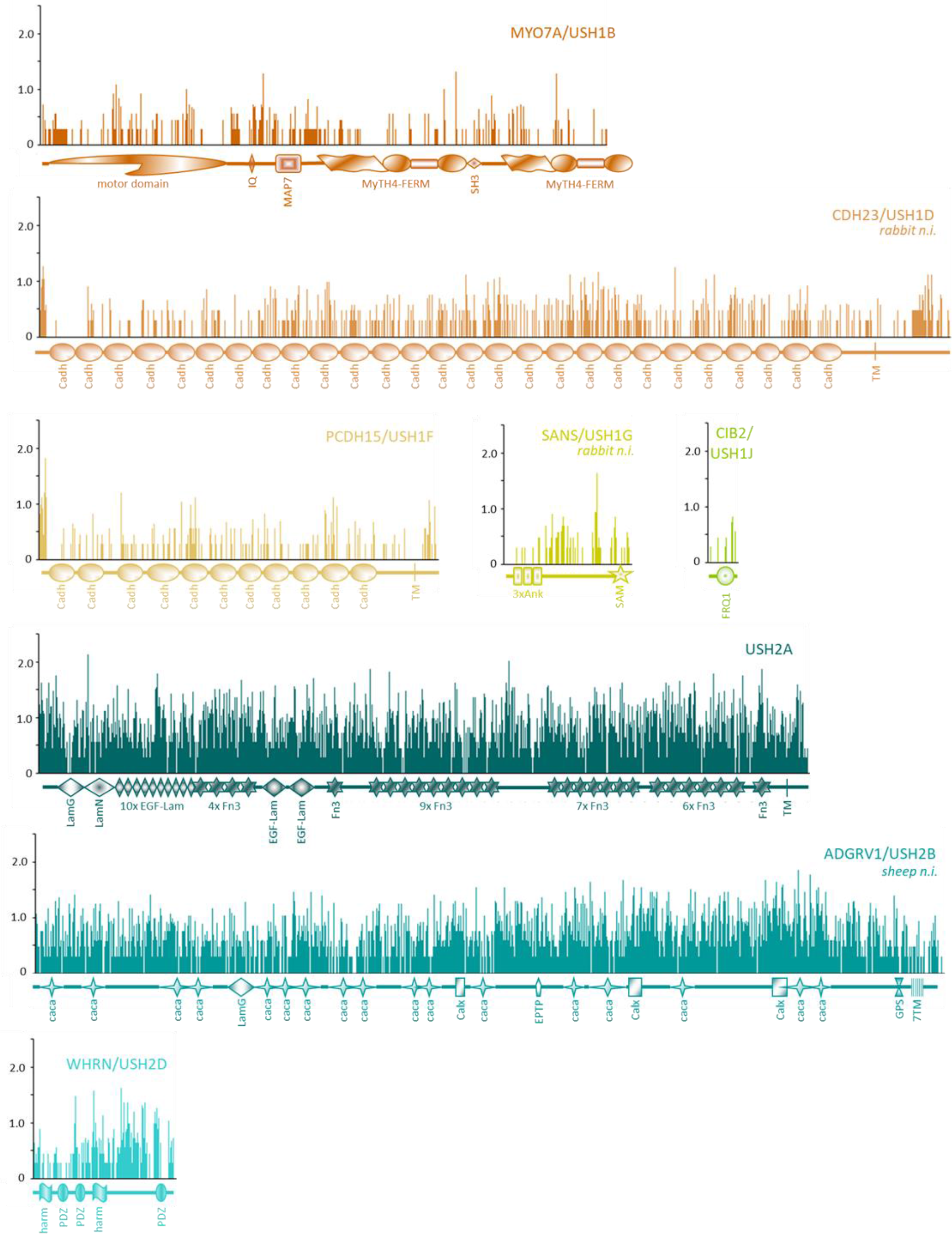
USH protein conservation. The protein sequences for USH1 and USH2 components from 12 species (human, macaque, marmoset, pig, cattle, sheep, horse, cat, dog, rabbit, mouse, rat) were fetched from the GenBank database, aligned and used for generation of Stewart entropy plots. If not available from the database, specific species were not included (n.i.) for given proteins. The principle structure of protein domains were taken from the respective protein descriptions in the GenBank entry or revealed by the search-for-conserved-domain algorithm (both at https://www.ncbi.nlm.nih.gov/) and indicated below the entropy plot. USH1 components appear substantially more conserved than USH2 components.

## Notes

### Competing Interest Statement

The authors have declared no competing interest.

## References

1. P. Mathur, J. Yang, Usher syndrome: hearing loss, retinal degeneration and associated abnormalities. Biochimica et Biophysica Acta (BBA)-Molecular Basis of Disease 1852, 406–420 (2015).

2. C. G. Moller et al., Usher syndrome: an otoneurologic study. Laryngoscope 99, 73–79 (1989).

3. M. Toms, W. Pagarkar, M. Moosajee, Usher syndrome: clinical features, molecular genetics and advancing therapeutics. Ther Adv Ophthalmol 12, 2515841420952194 (2020).

4. H. Lonborg-Moller, Y. Subhi, L. Kessel, Living with Usher Syndrome: Patient and Physician Perspectives. Ophthalmol Ther 9, 1–6 (2020).

5. A. El-Amraoui, C. Petit, The retinal phenotype of Usher syndrome: pathophysiological insights from animal models. C R Biol 337, 167–177 (2014).

6. D. S. Williams, Usher syndrome: animal models, retinal function of Usher proteins, and prospects for gene therapy. Vision Res 48, 433–441 (2008).

7. M. J. Chandler, P. J. Smith, D. A. Samuelson, E. O. MacKay, Photoreceptor density of the domestic pig retina. Vet Ophthalmol 2, 179–184 (1999).

8. A. Hendrickson, D. Hicks, Distribution and density of medium- and short-wavelength selective cones in the domestic pig retina. Exp Eye Res 74, 435–444 (2002).

9. H. May-Simera, K. Nagel-Wolfrum, U. Wolfrum, Cilia - The sensory antennae in the eye. Prog Retin Eye Res 60, 144–180 (2017).

10. C. Schietroma et al., Usher syndrome type 1–associated cadherins shape the photoreceptor outer segment. Journal of Cell Biology 216, 1849–1864 (2017).

11. I. Sahly et al., Localization of Usher 1 proteins to the photoreceptor calyceal processes, which are absent from mice. Journal of Cell Biology 199, 381–399 (2012).

12. U. Wolfrum et al., Subcellular Localization of Usher Syndrome Proteins in the Human Retina. Investigative Ophthalmology & Visual Science 51, 2494–2494 (2010).

13. S. Y. Li, Z. Q. Yin, S. J. Chen, L. F. Chen, Y. Liu, Rescue from light-induced retinal degeneration by human fetal retinal transplantation in minipigs. Curr Eye Res 34, 523–535 (2009).

14. J. Monés et al., A Swine Model of Selective Geographic Atrophy of Outer Retinal Layers Mimicking Atrophic AMD: A Phase I Escalating Dose of Subretinal Sodium Iodate. Invest Ophthalmol Vis Sci 57, 3974–3983 (2016).

15. W. Wang et al., Selective rod degeneration and partial cone inactivation characterize an iodoacetic acid model of Swine retinal degeneration. Invest Ophthalmol Vis Sci 52, 7917–7923 (2011).

16. C. Kostic et al., Rapid cohort generation and analysis of disease spectrum of large animal model of cone dystrophy. PLoS One 8, e71363 (2013).

17. R. M. Petters et al., Genetically engineered large animal model for studying cone photoreceptor survival and degeneration in retinitis pigmentosa. Nat Biotechnol 15, 965–970 (1997).

18. J. W. Ross et al., Generation of an inbred miniature pig model of retinitis pigmentosa. Investigative ophthalmology & visual science 53, 501–507 (2012).

19. J. R. Sommer et al., Production of *ELOVL4* transgenic pigs: a large animal model for Stargardt-like macular degeneration. British Journal of Ophthalmology 95, 1749–1754 (2011).

20. B. Boeda et al., Myosin VIIa, harmonin and cadherin 23, three Usher I gene products that cooperate to shape the sensory hair cell bundle. EMBO J 21, 6689–6699 (2002).

21. J. Reiners et al., Differential distribution of harmonin isoforms and their possible role in Usher-1 protein complexes in mammalian photoreceptor cells. Invest Ophthalmol Vis Sci 44, 5006–5015 (2003).

22. I. Zwaenepoel et al., Identification of three novel mutations in the USH1C gene and detection of thirty-one polymorphisms used for haplotype analysis. Hum Mutat 17, 34–41 (2001).

23. M. Kurome, B. Kessler, A. Wuensch, H. Nagashima, E. Wolf, Nuclear transfer and transgenesis in the pig. Methods Mol Biol 1222, 37–59 (2015).

24. P. Vochozkova, K. Simmet, E. M. Jemiller, A. Wunsch, N. Klymiuk, Gene Editing in Primary Cells of Cattle and Pig. Methods Mol Biol 1961, 271–289 (2019).

25. S. Ernest et al., Mariner is defective in myosin VIIA: a zebrafish model for human hereditary deafness. Hum Mol Genet 9, 2189–2196 (2000).

26. K. R. Johnson et al., Mouse models of USH1C and DFNB18: phenotypic and molecular analyses of two new spontaneous mutations of the Ush1c gene. Hum Mol Genet 12, 3075–3086 (2003).

27. S. Egerer et al., Early weaning completely eliminates porcine cytomegalovirus from a newly established pig donor facility for xenotransplantation. Xenotransplantation 25, e12449 (2018).

28. J. M. Dougherty, M. Carney, P. D. Emmady, in StatPearls. (Treasure Island (FL), 2020).

29. D. Bonneau et al., Usher syndrome type I associated with bronchiectasis and immotile nasal cilia in two brothers. J Med Genet 30, 253–254 (1993).

30. S. Pieke-Dahl et al., Genetic heterogeneity of Usher syndrome type II: localisation to chromosome 5q. J Med Genet 37, 256–262 (2000).

31. E. G. Puffenberger et al., Genetic mapping and exome sequencing identify variants associated with five novel diseases. PLoS One 7, e28936 (2012).

32. F. Barone et al., Behavioral Assessment of Vision in Pigs. Journal of the American Association for Labaratory Animal Science 57, 350–356 (2018).

33. D. Cvejic, T. A. Steinberg, M. S. Kent, A. Fischer, Unilateral and bilateral congenital sensorineural deafness in client-owned pure-breed white cats. J Vet Intern Med 23, 392–395 (2009).

34. G. Lefevre et al., A core cochlear phenotype in USH1 mouse mutants implicates fibrous links of the hair bundle in its cohesion, orientation and differential growth. Development 135, 1427–1437 (2008).

35. R. K. Koenekoop, M. A. Arriaga, K. M. Trzupek, J. J. Lentz, in GeneReviews((R)), M. P. Adam et al., Eds. (Seattle (WA), 1993).

36. D. S. Williams et al., Harmonin in the murine retina and the retinal phenotypes of Ush1c-mutant mice and human USH1C. Invest Ophthalmol Vis Sci 50, 3881–3889 (2009).

37. C. Hippert et al., Muller glia activation in response to inherited retinal degeneration is highly varied and disease-specific. PLoS One 10, e0120415 (2015).

38. A. Bringmann et al., Cellular signaling and factors involved in Muller cell gliosis: neuroprotective and detrimental effects. Prog Retin Eye Res 28, 423–451 (2009).

39. F. D. Gregory, T. Pangrsic, I. E. Calin-Jageman, T. Moser, A. Lee, Harmonin enhances voltage-dependent facilitation of Cav1.3 channels and synchronous exocytosis in mouse inner hair cells. J Physiol 591, 3253–3269 (2013).

40. F. S. Sjostrand, The ultrastructure of the outer segments of rods and cones of the eye as revealed by the electron microscope. J Cell Comp Physiol 42, 15–44 (1953).

41. L. S. Carvalho et al., Synthetic Adeno-Associated Viral Vector Efficiently Targets Mouse and Nonhuman Primate Retina In Vivo. Hum Gene Ther 29, 771–784 (2018).

42. L. H. Vandenberghe et al., AAV9 targets cone photoreceptors in the nonhuman primate retina. PLoS One 8, e53463 (2013).

43. M. Langin et al., Consistent success in life-supporting porcine cardiac xenotransplantation. Nature 564, 430–433 (2018).

44. A. Moretti et al., Somatic gene editing ameliorates skeletal and cardiac muscle failure in pig and human models of Duchenne muscular dystrophy. Nat Med 26, 207–214 (2020).

45. A. P. Regensburger et al., Detection of collagens by multispectral optoacoustic tomography as an imaging biomarker for Duchenne muscular dystrophy. Nat Med 25, 1905–1915 (2019).

46. I. Ebermann et al., PDZD7 is a modifier of retinal disease and a contributor to digenic Usher syndrome. J Clin Invest 120, 1812–1823 (2010).

47. J. C. Corral-Serrano et al., PCARE and WASF3 regulate ciliary F-actin assembly that is required for the initiation of photoreceptor outer segment disk formation. Proc Natl Acad Sci U S A 117, 9922–9931 (2020).

48. W. J. Spencer, T. R. Lewis, J. N. Pearring, V. Y. Arshavsky, Photoreceptor Discs: Built Like Ectosomes. Trends Cell Biol, (2020).

49. U. Wolfrum, A. Schmitt, Rhodopsin transport in the membrane of the connecting cilium of mammalian photoreceptor cells. Cell Motil Cytoskeleton 46, 95–107 (2000).

50. P. Kiesel et al., The molecular structure of mammalian primary cilia revealed by cryo-electron tomography. Nat Struct Mol Biol 27, 1115–1124 (2020).

51. J. Kim et al., Actin remodelling factors control ciliogenesis by regulating YAP/TAZ activity and vesicle trafficking. Nat Commun 6, 6781 (2015).

52. A. Karam et al., A novel function of Huntingtin in the cilium and retinal ciliopathy in Huntington’s disease mice. Neurobiology of Disease 80, 15–28 (2015).

53. K. M. Bujakowska, Q. Liu, E. A. Pierce, Photoreceptor Cilia and Retinal Ciliopathies. Cold Spring Harb Perspect Biol 9, (2017).

54. S. Khateb et al., Phenotypic Characteristics of Rod-Cone Dystrophy Associated with Myo7a Mutations in a Large French Cohort. Retina 40, 1603–1615 (2020).

55. K. Stingl et al., Full-field electroretinography, visual acuity and visual fields in Usher syndrome: a multicentre European study. Doc Ophthalmol 139, 151–160 (2019).

56. E. Lenassi et al., Natural history and retinal structure in patients with Usher syndrome type 1 owing to MYO7A mutation. Ophthalmology 121, 580–587 (2014).

57. S. Veleri et al., Biology and therapy of inherited retinal degenerative disease: insights from mouse models. Dis Model Mech 8, 109–129 (2015).

58. S. Russell et al., Efficacy and safety of voretigene neparvovec (AAV2-hRPE65v2) in patients with RPE65-mediated inherited retinal dystrophy: a randomised, controlled, open-label, phase 3 trial. Lancet 390, 849–860 (2017).

59. C. Kostic, Y. Arsenijevic, Animal modelling for inherited central vision loss. J Pathol 238, 300–310 (2016).

60. A. J. Zele, D. Cao, Vision under mesopic and scotopic illumination. Front Psychol 5, 1594 (2014).

61. M. Karlstetter et al., Disruption of the retinitis pigmentosa 28 gene Fam161a in mice affects photoreceptor ciliary structure and leads to progressive retinal degeneration. Hum Mol Genet 23, 5197–5210 (2014).

62. M. Dominik Fischer et al., Detailed functional and structural characterization of a macular lesion in a rhesus macaque. Doc Ophthalmol 125, 179–194 (2012).

