## Supplemental Methods for "Early Disruption of Photoreceptor Cell Architecture and Loss of Vision in a Humanized Pig Model of Usher Syndrome"

***Supplemental materials & methods***

### Establishing the pig model

Modification of the porcine *USH1C* locus was carried according Vochozkova et al. (Vochozkova et al., 2019) and generation of genetically modified pigs by somatic cell nuclear transfer (SCNT) was done according Kurome et al (Kurome et al., 2015) (Figure 2a). The following aspects were specific for the generation of the USH pig model (for details see the oligo & gene synthesis section below):

Bioinformatics evaluation: *USH1C* loci were identified in macaque, marmoset, cattle, sheep, pig, horse, cat, dog, rabbit, mouse, and rat by BLAT using human exonic sequences and extracted from the respective reference genomes (www.ensembl.org). Multi-species alignments were done by a combination of DiAlign and CHAOS (Brudno et al., 2004). Proposed regulatory elements were extracted from the USCS Genome browser (genome.ucsc.edu) as DNase sensitive elements (Thurman et al., 2012), ENCODE ChIP-Chip (Wang et al., 2013), FANTOM5 enhancer elements (Andersson et al., 2014), GeneHancer elements (Fishilevich et al., 2017), FAIRE ENCODE regions (Giresi et al., 2007) and PreMod (Ferretti et al., 2007). Any identified regulatory element was localized within the alignment in BioEdit (https://bioedit.software.informer.com/7.2/) and a density plot was created. Both, a homology plot at nucleic acid level and a protein conservation plot were calculated in BioEdit on the basis of the nucleotide summary assessement and entropy calculation functions, respectively. Coverage maps of the augmented alignment were plotted by JalView (Waterhouse et al., 2009).

Personalized R31X allele: For introducing a patient-specific segment, we analyzed a 1.6kb fragment from an individual carrying a USH1C$^{R31X}$ mutation (Figure 1b). The PCR product was amplified with the primers *hush1c_2f* and *hush1c_2r* (for details see the oligo & gene synthesis section below), cloned into a plasmid vector and sequenced. For avoiding the misinterpretation of potential PCR mistakes, the sequence of the R31X-allele was based on 6 distinct clones and the other allele on 3 distinct clones. Consistently appearing polymorphisms were defined as SNPs.

Targeting construct: The porcine *USH1C* gene has been annotated to chr2:42Mb, with the exons largely corresponding to the human *USH1C* transcript variant b3, GenBank no. NM_153676.4 (Figure S1). Different porcine primary cell lines (Richter et al., 2012) were examined for SNPs in the target region and 3 distinct gRNAs, *rk1*, *rk3* and *rk4* that were not affected by these SNPs were examined for their power to introduce NHEJ-based mutations in pig primary cells lines; for this, cells were nucleofected and after 48 hours, the target locus amplified with the primers *ushwt1f* and *ushwt1r*, cloned into plasmids and sequenced. Targeting experiments were based on *rk4* as it proved superior (12% mutation rate) to *rk1* (6%) and *rk3* (10%). BACs CH242-515C3 and CH242-199J14, were identified to completely carry the porcine *USH1C* locus via the Sscrofa 10.2 reference genome and the PigPre BAC map (www.ensemlb.org) (Figure S3a) and purchased from CHORI BACPAC Resources Center (BACPAC Genomics, Emeryville, CA, USA). A sequence comprising a 278bp 5´-arm, the 1524bp humanized fragment including the TGA stop codon in exon 2 and 3 intronic SNPs as well as a 278bp 3´-arm were commercially synthesized and provided in a pUC plasmid derivate by GeneArt (Invitrogen by Thermo Fisher Scientific, Carlsbad, CA, USA). A cassette with a PGK/EM7 driven *neo* gene, providing neomycin resistance in mammalian cells and kanamycin resistence in bacteria (Klymiuk et al., 2012), was cloned into the plasmid via the *Not*I/*Xho*I sites located between the 5´-arm and the humanized fragment. The entire fragment was excised from the backbone via *Asc*I and introduced into the CH242-515C3 BAC by bacterial recombineering, according Vochozkova et al (Vochozkova et al., 2019) using the SW106 E.coli strain (Warming et al., 2005). Correct modification of the BAC was proven by PCR for recombination at the 5´- and 3´-ends with the primer pairs *5arm1f – 5arm1r* and *3armwf – 3arm2r,* and for the removal of the corresponding porcine segment with the primer pair *ushwt1f – ushwt1r*. Further, integrity of the BAC and changes in the RE pattern due to the modification were confirmed by digestion with *Xba*I and *Spe*I (Figure S3b).

Generating pigs: Plasmids carrying the modified BAC, the gRNA *rk4* and Cas9 were prepared endotoxin-freely and transfected into pig primary cells PKCf and PKCm by nucleofection (Amaxa by Lonza, Basel, Switzerland). Single cell clones were generated as described (Richter et al., 2012) and propagated towards a 2x 96-well scale. One aliquot was used for analysis by the loss-of-wildtype-allele approach (LOWA) (Vochozkova et al., 2019) and the other served as a backup for potential SCNT. qPCR was carried out on a LightCycler96 (Roche Life Science, Basel, Switzerland) using FastStart Essential DNA Green Master (Roche Life Science) and primer pair *ush1c_qf1 – ush1c_qr1* as well as the primer pair *o4_qf2 – o4_qr1* for a reference site in the POU5F1 gene and *ng_qf6 – ng_qr4* for a second reference in the NANOG gene. Verified single cell clones were used as donors in SCNT and embryos were transferred to synchronized gilts, according standard procedures (Kurome et al., 2015). Birth was introduced in pregnant foster mothers by Estrumate at day 115. (MSD Animal Health, Merck, Kenilworth, NJ; USA).

Tissue sampling: For isolation of pig primary kidney cells and sampling of tissue, animals were sedated with ketamine 100mg/ml (Ursotamin®, Serumwerk Bernburg, Germany) and azaperone 40mg/ml (Stresnil®, Elanco Animal Health, Bad Homburg, Germany), according to the manufacturer's specifications. Fully anesthetized animals were euthanized by intravenous injection of T61® (MSD Animal Health). Pig primary kidney cells were isolated according to standard procedures (Richter et al., 2012) (see below). Tissue samples for histology, electron microscopy and molecular analyses were collected according to established sampling guidelines adapted to porcine biomedical models (Albl et al., 2016). Samples for molecular analysis were frozen on dry ice and stored at -80°C. Tissue was powdered using hammer and anvil first and pestle and mortar afterwards. All instruments and samples were cooled in liquid nitrogen to avoid thawing of the samples during processing. Powdered tissue was transferred to a pre-cooled tube and stored at -80°C.

Breeding: After reaching fertility and stabilization of cycle, USH1C F0 sows were inseminated with WT sperm. When pregnant, sows were trained and reassured intensively for farrowing by regular physical contact with caretaker. F1 offspring were genotyped (see below) and raised for breeding purposes. After reaching fertility F1 boars were mated to their mothers and sisters for producing USH1C KO pigs and with wild-type sows to broaden the genetic background of the future breeding herd. Litters from hom x het matings were with their mother for suckling, but piglets kept in a separate box otherwise. Upon calling by the mother, the barrier between the boxes was opened and the time until piglets passed the gate was determined. Non-responding piglets were nudged by the caretaker after one minute to avoid impaired nurturing.

**Molecular analysis**

Primer sequences are summarized in the oligo & gene synthesis section (see below).

Genomic level: Genomic DNA was isolated from tail biopsies by using the Easy DNA kit (Invitrogen) DNAeasy kit (Qiagen, Hildesheim, Germany) or Nexttec kit (Nexttec GmbH, Leverkusen, Germany). For verifying the abundance of the modification element, amplicons generated with the primer pair *hush_for3 – hush_rev3* were sequenced. The presence of the neo selection cassette was detected by PCR with *usharm2f – ush5arm2r* and sequencing whereas Cre-mediated excision of neo was detected by PCR with *neoFOR3 – neoREV3* and sequencing.

Transcriptional level: RNA was isolated from powdered tissue samples using TRIzol™ (Invitrogen). Approximately 100mg of powdered tissue was grinded up in 1ml of Trizol by Polytron PT2500E (Kinematica, Luzern, Switzerland) in a pulsatile manner to avoid overheating. Further steps were carried out as suggested by the Trizol protocol and RNA was stored at -80°C. For cDNA synthesis, samples were treated first with DNase I (Invitrogen) to remove a possible contamination with gDNA. Then, RNA was reversely transcribed into cDNA with SuperScript™ III Reverse Transcriptase (Invitrogen). For determining *USH1C* transcripts, cDNA was used in RT-PCRs with the primer pair *ushrt2f – ushrt2r* or *ushrt2f – urt4r*, located in the consistently transcribed exons 1 and 8 or 6, respectively. Afterwards, Sanger sequencing was performed on PCR amplicons.

Protein analysis: Tissue was powdered and then processed as previously described (Sedmak and Wolfrum, 2011). In brief, retinal tissues were lysed in modified RIPA buffer (50 mM Tris–HCl, 150 mM NaCl, 0.1% SDS, 2 mM EDTA, 1% NP-40, 0.5% sodium-deoxycholate, 1 mM sodiumvanadate, 30 mM sodium-pyrophosphate, pH 7.4). Protein lysates were separated by SDS-PAGE gel electrophoresis, followed by semi-dry Western blotting as previously described in (Overlack et al., 2011). Western blots were analyzed with the Odyssey infra-red imaging system (LI-COR Biosciences, Lincoln, NE, USA). For details on antibodies see below.

**Morphology & Histology**

Immunohistochemistry: Porcine eyes were fixed in melting isopentane for cryosectioning as described in (Overlack et al., 2011). Samples were sectioned with a MICROM HM 560 Cryo-Star cryostat (Fisher Scientific by Thermo Fisher Scientific, Waltham, MA, USA) and placed on poly-L-lysine-precoated coverslips. After drying, samples were incubated with 0.01% Tween 20 PBS, then washed with PBS and covered with blocking solution (0.5% cold-water fish gelatin, 0.1% ovalbumin in PBS) for a minimum of 30 min followed by over-night incubation at 4 °C with primary antibodies (for details on antibodies and fluorescent dyes see below). For GFAP staining samples were postfixed with 4% paraformaldehyde (PFA) in PBS. Cryosections were washed several times and incubated with secondary antibodies in blocking solution containing DAPI (Sigma-Aldrich, St Louis, MO, USA) for 1 h at room temperature. After washing, sections were mounted in Mowiol (Roth, Karlsruhe, Germany). Slides were analyzed on a Leica DM6000B microscope (Leica, Bensheim, Germany); images were processed with LAS-AF Leica imaging software and ImageJ/Fiji software (Schindelin et al., 2015) or Adobe Photoshop CS (Adobe Systems, San Jose, CA, USA).

Scanning electron microscopy (SEM): The temporal bones of the pigs were dissected to expose the middle and inner ears. Their cochleae were dissected out immediately and the organ of Corti was exposed. The tectorial membrane was removed prior to fixation. Dissected cochleae were fixed (in 2,5 % glutaraldehyde, 4% PFA in 0,1 M Sörensen's phosphate buffer). After several washing steps with 0,1 M Sörensen's phosphate buffer, cochleae were dehydrated in an ethanol series, critical-point dried and gold sputtered in an argon athmosphere. Specimen were imaged with a Philips ESEM XL30 scanning electron microscope (Philips, Eindhoven, Netherlands).

Transmission electron microscopy (TEM): For conventional TEM dissected eyeballs were pre-fixed for 2 hours in buffered 2.5% glutaraldehyde containing sucrose and post-fixed in buffered 2% OsO4 as previously described (Karlstetter et al., 2014). After dehydration in ethanol series and the passage through propylenoxid as an intermedium, samples were embedded in Renlam® M-1 and polymerized at 60°C. For the pre-embedding labeling we followed a previously established protocol (Sedmak and Wolfrum, 2010). In brief, eyes were pre-fixed in buffered 4% PFA, dissected, infiltrated with 30% buffered sucrose, and cracked by freezing - thawing cycles, followed by embedding in buffered 2% Agar (Sigma-Aldrich). Agar blocks were sliced with a VT1000 S vibratome (Leica). Endogenous peroxidase activity of vibratome sections was suppressed by incubation with $H_2O_2$. Sections were incubated with primary antibodies for 4 days and overnight with biotinylated secondary antibody and were visualized with a Vectastain ABC-Kit (Vector Laboratories, Burlingame, CA, USA).

Retina sections were post-fixed first in buffered 2.5% glutaraldehyde and second in buffered 0.5% OsO4. After dehydration, sections were flatmounted between ACLAR®-films (Ted Pella Inc., Redding, CA, USA) in Renlam® M-1 resin. Flatmount samples were heat-polymerized and glued on top of empty Araldit blocks. Ultrathin sections of the embedded specimens were prepared with an Ultracut S ultramicrotome (Leica) and collected on Formvar-coated copper or nickel grids. Sections were counter stained with heavy metals before being analyzed and imaged in a Tecnai 12 BioTwin TEM (FEI company by Thermo Fisher Scientific, Hillsboro, OR, USA) equipped with a SIS Mega-View3 CCD camera (EMSIS, Münster, Germany) was used. The images were processed using Adobe Photoshop CS (Adobe Systems). Measurements of structures were done using the analysis system of SIS. Quantification steps have been performed with Fiji/ImageJ. The primary ciliary length of fibroblasts was measured by hand. For the analysis of retinal tissue, including length measurements of connecting cilia and width measurements of cone pedicle synapses, macros were processed in Fiji/ImageJ for an automatic evaluation of the fluorescent pictures.

**Primary cell culture:**

Pig primary kidney cells and primary dermal fibroblasts were isolated according to standard procedures (Richter et al., 2012). In brief, kidneys were taken from euthanized animals, washed in PBS containing Penicillin/Strepomycin and minced. Tissue was treated with collagenase and filtered cells were cultivated in DMEM. Pieces of porcine skin were disinfected in 98% Ethanol and after short washing in PBS placed in Betaisodona 10%-solution. After several washing steps in PBS, skin was minced. Kidney or skin samples were then completely covered with MEM containing 10% heat-inactivated fetal calf serum (FCS) and 1% penicillin-streptomycin (PS) and incubated at 37°C and 5% CO2. 5 days later 10% FCS were added to the skin. Outgrown cells were transferred to new complete medium for further culturing.

For ciliogenesis experiments, $5x10^5$ cells were seeded per well (in medium containing 10 % FCS and 5 % PS) and 24 h later cultured in OPTI-MEM reduced-serum medium (Invitrogen by Thermo Fisher Scientific). After 48 h of starvation, cells were washed in PBS and fixed with 2% PFA in PBS. After washing and permeabilization with 0.1% Triton-X in PBS, cells were blocked with 0.5% cold-water fish gelatin, 0.1% ovalbumin in PBS followed by incubation with primary antibodies overnight at 4°C. After washing, cover slips were incubated with secondary antibodies for 1 h at room temperature. Samples were washed and then mounted in with Mowiol (Roth).

**Clinical examination**

Anesthesia: For examination with ERG, mfERG, OCT, AF, FA, and ABR on 3-week old animals were anesthetized by intramuscular injection of 10 mg/kg azaperon (Stresnil®, Elanco, Germany), 0.02 mg/kg atropine sulfate (B. Braun, Germany) and 20 mg/kg ketamine (Ursotamin®, Serumwerke Bernburg, Germany). Anaesthesia was continued by intravenous injection of propofol (Fresenius, Germany) according to effect. After endotracheal intubation, pigs were mechanically ventilated and cardiovascular function was monitored throughout the procedures. To exclude eyeball movement during examinations, a peripheral muscle relaxant (Rocuronium, Inresa, Germany) was applied. For ABR on older animals, animals were starved over night befor intramuscular application of TKX (tiletamine 4 mg/kg, zolazepam 4 mg/kg (Zoletil 100; Virbac), ketamine 5 mg/kg (Narketan 10; Chassot), and xylazine 1 mg/kg (Rometar 2%; Spofa).

Auditory Brain Stem Response (ABR): In 3-week-old USH1C and WT piglets, ABR was recorded with standard electrodiagnostic equipment (Viking Quest®; Natus, Planegg, Germany) and Natus® TIP-300 insert earphones (AF). Recording stainless steel needle electrodes were positioned subcutaneously ipsilateral to the stimulated ear over the mastoid at the base of the ear (-) and at the vertex in the midline of the skull corresponding to Cz (+). A ground electrode was positioned in the neck. Electrode impedance was < 2kOhm. Click stimuli (100µsec, alternating polarity, 11.1 Hz) were delivered at supramaximal stimulation intensity (100 dBSPL) to one ear and masking noise was applied to the contralateral ear (-40 dBSPL). Filter settings were 100 Hz for high pass and 3 kHz for low pass filters. The ABR represented the averaged signal of 1000 - 1500 10 msec recordings. ABR was recorded twice for each ear to ensure reproducibility of the ABR peaks in WT animals (n = 3) or absence of recognizable peaks in $Ush^{-/-}$ animals (n = 3), respectively. If there was no recordable ABR at 100 dBSPL in $Ush^{-/-}$ animals, ABR was repeated at 120 dBSPL (twice for each ear). Additional testing comprised ABR at 10 dB decreasing steps in WT piglets. Hearing threshold was determined as the minimal click intensity that still evoked a noticeable potential negative peak after peak V.

ABR for other timepoints were conducted with four subcutaneous needles placed in vertex, forehead and mastoids with spezialized audiometric equipment. 8 weeks and 2 years old animals were examined with eABRUSB (eABR – BioMed Jena GmbH, calibrated by standard norm IEC 318-4) and headphones (Sennheiser Momentum M2 In-Ear G Black-Red, 18Ω) were used on the 8 weeks old piglets. Click stimuli (duration 100us, steps 10 to 5 dB) and tone pips (duration 100us, frequencies 2, 4, 8, 16 kHz, steps 10 to 5 dB) were applied. The evoked signal was filtered by band-pass filter of range from 300Hz to 3kHz. All the responses were averaged 200-400 times for each intensity and analysed with eAudio software. On 3 months, 6 months and 2 years old pigs, acoustic stimuli were generated with an evostar 2 system (ERA system: Evoselect/Evostar 2, Pilot Blankenfelde medizinisch elektronische Geräte GmbH) and presented to the animal via headphones (beyerdynamic, DT48, 5Ω, Germany). Click stimuli (duration of electric pulse 150 us) and the tone pips (duration 2 ms, frequencies of 0.5, 1, 2, 3, 4 kHz, intensity steps from 10 to 5 dB) were applied. The signal from an electrode was filtered by band-pass filter over the range of 100Hz to 2,5kHz. The response was averaged 200-500 times on each intensity. This signal was processed with and analyzed using Evostar software. The threshold response to each frequency and click was determined as the minimal tone/click intensity that still evoked a noticeable potential peak in the expected time window of the recorded signal. This evaluation process was the same for both systems.

<u>Electroretinography (ERG):</u> Full-field ERG was recorded with a corneal Kooijman electrode (Roland Consult, Brandenburg an der Havel, Germany). Platinum needles were placed subcutaneously at the temple and top of the head between the ears as reference and ground electrodes respectively. Stimuli were brief white flashes (4 ms) delivered from a light source within the Kooijman electrode (Kooijman and Damhof, 1986). The RETImap system (Roland Consult) was used for stimulus generation and data acquisition. The recording protocol used was based on, and extended from, the International Society for Clinical Electrophysiology of Vision standard for human clinical full-field ERG. A band pass filter was used to remove frequencies outside 1 - 300Hz. An interstimulus interval (ISI) of 1 s was used with 20 responses recorded and averaged for each single flash protocol. For scotopic ERG, the protocol consisted of 30 minutes dark-adaptation, followed by recording of a dark-adapted single flash series with increasing luminance from −3.0 log cd*s*m−2 to 2.0 log cd*s*m−2. After completion of the dark-adapted intensity series, animals were light adapted for 10 minutes to a steady white background of 30 cd*s*m−2 with the same light source. Light-adapted standard flash was recorded by using white flash stimuli at 0 log cd*s*m−2. The same luminance was used to record a 28Hz flicker response for 300ms. For multifocal ERG, the mfERG was recorded using the same RETImap system (Roland Consult) using a 61-hexagon stimulus and according to the standard of the International Society for Clinical Electrophysiology of Vision (Hood et al., 2012). A 61-hexagon stimulus was presented on a 51-cm cathode ray tube monitor with a frame frequency of 60 Hz. It encompassed the central visual field of 44° horizontally and vertically. The scaled-size hexagons (distortion factor 4) were light-modulated according to a binary pseudorandom m-sequence. The light state had a luminance of 120 cd/m−2 and the dark state of 1 cd/m−2. Refractive errors were not corrected. Pupil dilation was achieved with one drop of tropicamide–phenylephrine mixture into the conjunctival sac of each eye about 15 min before the exam. Contact lens electrodes (ERG-jet®, CareFusion, San Diego, USA) were applied. Amplifier gain was set to 50,000 and the band pass from 5 to 100 Hz. There were eight recording cycles of 47 s each. The 61 traces are displayed separately in a false-colour coded amplitude map identifying P1 amplitude (measured from the N1 trough to the following maximum positive peak; N1 amplitude was measured from the electroneutral starting point to the first negative deflection). All traces were also grouped into one response value and the averaged first-order kernel function displayed as single trace. Luminance levels in the pig eye were calculated from the light intensity measured at the floor. Given the complex physical correlation between light source, reflection and sensing in an indoor 3D space, it is challenging to determine corresponding luminance levels in the pig eye, but considering the general reflexion parameter $\rho$ as 0.2 seems a fair approximation.

<u>Optical Coherence Tomography (OCT):</u> Spectral-domain OCT B-scans of central retina in animals were obtained using a Spectralis HRA+OCT device (Heidelberg Engineering, Heidelberg, Germany) under general anaesthesia and during full pupil dilation. During the acquisition, the corneal surface was protected with methylcellulose eye drops while lids were held open using lid specula. After 30deg and 55deg infrared and red-free images were recorded using scanning laser ophthalmoscopy (SLO), animal received a flush of approximately 5 mg/kg fluorescein and the angiography was recorded using the 55 degree lens at multiple time points (0.5, 1, and 5 minutes).

**Behavior testing**

Barrier course:

This course was adapted from a previous publication (Barone et al., 2018). Ten barriers (boards of 94x76cm) were placed in a distance of one meter in a straight aisle (12x2.2m) alternating in the middle, the left and right. There were four different arrangements of barriers, which were used alternatingly (Fig S2) Pigs were trained in the course until they walked straight through the course without pausing or hesitating for at least two times in a row. After a training period of four weeks, the animals were tested in the course weekly. The pigs entered the course individually in random order, they had to pass the barriers and at the end they were rewarded with fruit juice in a food bowl. The test was performed before feeding to ensure a higher motivation in the pigs. After some runs in the light, the test was conducted alternately in the light (average of 135 lux) and in the dark (average of 2.9 lux). Illumination level was measured at three different sites in the course (start, middle and end) in animals' head height.

Each run was documented by video. The pigs were marked with numbers on their backs to facilitate recognition. Times the pigs took to pass the course, from snout crossing the start line until the pig was touching the food bowl, were measured. All barrier contacts by the pigs were counted. Contacts were divided into two subgroups. Frontal contacts mean the pig was touching the barrier with its snout or head, when the pig was touching the obstacle barrier with its shoulders or hips, the contacts were called lateral contacts.

Each time a pig was turning around and walking back before reaching the end was counted. It was distinguished between turns in front of the course and turns in the course. Such runs were excluded. After turning around the animals were sent back in the course, when they walked to the end then, this new run was documented with time and contacts. Also every time a pig was walking in circles was counted. As long as they were just circling and not walking back, the run was not excluded, time measurement was continuing. When the pig was circling before entering the course (that means in front of the start line), those circles were also counted, but time measurement only started when the animal was crossing the start line.

Pigs were excluded from one run when they were lame or when a pig had already been outside of its pen on the barrier course day for another reason. When a pig was turning around or circling so often that it needed longer than two minutes to walk to the endpoint, it was also excluded from this day's run. The numbers of turns/circles within those two minutes were still counted. Pigs were not included in the study due to lack of motivation (the pig did not go to the endpoint reliably after six weeks of training or was more interested in playing with the barriers than walking straight to the food bowl).

Obstacle course:

An obstacle course to test the visual capacity of USH1C pigs and WT pigs was conducted. The course was 6.15m long, the width was 1.58m in the beginning and 2.48m in the end. Most obstacles were built from obstacles used for equestrian sport, other obstacles were a stepboard and barrels. Obstacles were rearranged after almost all test days (Fig S3).

Pigs were trained until they walked straight through the obstacle course to the food trough without stopping or hesitating. After a training period of three weeks, the pigs were performing the course once a week. The pigs entered the course individually in random order, they had to pass the obstacles and at the end they were rewarded with a cookie in a food trough. The test was conducted under light and dim conditions.

Each run was documented by video. The pigs were marked with numbers to facilitate recognition. Times the pigs needed to pass the course were measured. All contacts with the obstacles or the wall were counted. When a pig nudged one obstacle more than once in a playful way in the same area, this was still counted as one touch. When a pig touched one obstacle at different sites, each contact was counted separately. When a pig stroked along one obstacle with its snout, this was counted as two touches. When a pig turned around before reaching the end or when a pig was lame, the pig was excluded from this test day.

A map of the obstacle course was drawn to see which route the pigs took through the course (Macromedia FreeHand MX, STUDIOMX 2004). Therefore, the dimensions of the course and the gaps of the slatted floor were measured and a plan was designed to a scale of one to ten. The videos of each animal and test day were watched and every step of the left and right forelimb was recorded in the map. The X and Y coordinate of each footstep with the corresponding time was then transferred to an Excel file. Trajectories were analyzed with the R package trajr. Briefly, we first imported and then smoothed all the trajectories to avoid any source of noise by applying a Savitzky-Golay smoothing filter (Luo et al., 2005; Savitzky and Golay, 1964). Then, we called all the functions of the trajr packages (McLean and Skowron Volponi, 2018) to characterize the trajectories into measures of speed, length, distance, duration and measures of straightness or tortuosity. The trajectory significance was tested using Kruskal-Wallis test.

For statistical evaluation, data were checked for consistency and normality. Fisher's Exact test or Pearson's test were used to analyze cross tabulations. Generalized linear models with Poission distribution, Median tests, bootstrap-t tests based on 5000 Monte Carlo simulations, t-tests with and without the assumption of homogeneity, Mann-Whitney U tests were used to test continuously distributed variables. All reported tests were two-sided, and p-values < 0.05 were considered statistically significant. Analyses were performed by use of NCSS_10 (NCSS LLC, Kaysville, UT, USA), STATISTICA_13 (Statsoft, Tulsa, OK, USA), MATHEMATICA_12 (Wolfram Research, Hanborough, UK), Champaign, IL (2018) and PASW 24 (SPSS by IBM Corp, Armonk, NY, USA) and Prism_5 (GraphPad Software, San Diego CA, USA).

**Therapy approaches:**

AAV-application: AAVs (AAV8.CMV.eGFP, AAV9.CMV.eGFPand Anc80.CMV.eGFP) were generated by the viral core facility at Boston Children's Hospital and the Gene Transfer Vector Core at the Massachusetts Eye and Ear Infirmary (Pan et al., 2017). AAVs were subretinal injected into the eyes of WT pig at an age of 30 months using an ophthalmologic surgery microscope (Hi-R NEO 900A, Haag-Streit). Using inhalation anesthesia (Morpheus E, Siare), —morphology of the retina was examined via OCT iVue (Optovue, Fremont, CA, USA) on both eyes before the injection. AAVs were applied in the subretinal space with Extendible 41G subretinal injection needle (23 gauge / 0.6 mm) (DORC, Zuidland, Netherlands). After 5 weeks injection sites were evaluated by OCT (see above) and funduscopy, confirming the concise injection sites. Following enucleation, the eye cup was dissect and fixed with 4% PFA in PBS. Flattened whole mount preparations of the eye cup were analyzed by epifluorescence using an handhold device (Bluestar Flashlight, Nightsea, Lexington, MA, USA). Samples were processed for cryosecting, cryosectioned and analyzed were analyzed on a Leica DM6000B microscope as described above.

Gene therapy: For restoring gene expression, 2.5x105 primary porcine fibroblasts were transfected with 0.4 µg of an endotoxin-free pDest_Harm_a1_S/F plasmid (Invitrogene), expressing splice form a1 of the human *USH1C* gene under the control of the CMV promoter. Transfection was performed with the 4D-NucleofectorTM X Unit from Lonza, using the P2 Primary Cell 4D-NucleofectorTM X Kit S and program FS113, according to the manufacturer's protocol. After transfection cells were seeded (in medium containing 10% FCS) and analyzed for ciliogenesis as described above.

Gene Repair: Different constellation of the gRNAs *urg1* and *urg2* with correcting ssODN *urt1*, *urt2*, *urt1.1*, *urt1.2*, *urt1.3*, *urt1.4* and *urt1.5* were tested in primary fibroblasts. For this, plasmids encoding Cas9 and the respective gRNA as well as correcting oligonucleotides were nucleofected under standard conditions (Richter et al., 2012). After 24-48 hours of cultivation, DNA was isolated from mixed cell batches and the efficacy of NHEJ and HDR were determined by examination of Sanger electropherograms of amplicons generated by PCR with the primer pair *huUSH2f* – *huUSH2r* and sequenced with primer ush1s in ICE CRISPR Analysis Tool (Synthego, Menlo Park, CA, USA). Cells nucleofected with *urg1* and *urt1.3* were seeded onto 96-well plates to generate single cell clones. For each clone identified, DNA was isolated, PCR amplicon was generated by primer pair *ushHS1f* – *huUSH1r*, sequenced with primer *ush1s* and analyzed by Sanger sequencing.

**Antibodies & Fluorescent dyes:**

The following primary antibodies were used in this study:

- rabbit polyclonal antibody against GFAP (#ZO334, purchased from DAKO by Agilent, Santa Clara, CA, USA); WB 1:10000, IF 1:1000
- affinity-purified rabbit polyclonal antibody against harmonin (H3) (homemade as in (Reiners et al., 2003)); WB 1:1000, IF 1:500
- rabbit polyclonal anti-Arl13b (#PTG 17711-1-AP, purchased from Acris by Origene, Rockville, MD, USA); IF 1:400
- goat polyclonal anti-pericentrin 2 (#sc-28145, purchased from Santa Cruz Biotechnology, Dallas TX, USA); IF 1:200
- mouse monoclonal antibody against Cent3 (homemade as in (Trojan et al., 2008); IF 1:100
- guinea-pig polyclonal anti-whirlin (homemade as in (van Wijk et al., 2006)); WB 1:5000
- rabbit polyclonal antibody against SANS (homemade as in (Sorusch et al., 2017)); WB 1:500
- rabbit polyclonal anti-myosinVIIa (#PTS-25-6790-CO50, purchased from Axxora by Enzo Life Sciences, Farmingdale, NY, USA); WB 1:1000
- mouse monoclonal antibody against GT335 (AdipoGen Life Sciences, (SanDiego, CA, USA); IF 1:1000
- mouse monoclonal anti-actin (#Sct MA5-11869, purchased from Invitrogen); WB 1:2000

Secondary antibodies used in this study were conjugated to Alexa488, Alexa 555, Alexa568, or Alexa 647, purchased from Invitrogen or Rockland Immunochemicals (Pottstown, PA; USA). For staining of cone pedicle synapses, lectin PNA (Peanut agglutinin) was coupled to Alexa 568. Nuclear DNA was stained with DAPI (4',6-diamidino-2-phenylindole) (1 mg/ml) (Sigma-Aldrich).

**Oligo & Gene Synthesis:**

Primers (with RE recognition sites underlined):

Genotyping:

- hush1c_2f: 5´-CTTAGGATCCAGGGAAGGAGGTACTGTCAGATCTAGG-3´
- hush1c_2r: 5´-TAAGCTCGAGACCACAAGTTACTCAAGGAAAGA-3´
- ushwt1f: 5'-TTCCCTCCTCTTCTCACAACCAT-3'
- ushwt1r: 5'-AGAGCCACGCACAGCACCTCT-3'
- 5arm1f: 5´-GGCAAGCTGTCCAGGTAGA-3´
- 5arm1r: 5´-TGCTTCACTGCTCCCAGAT-3´
- 3arm2f: 5´-TGTTGCCGTGAGCTGTG-3´
- 3arm2r: 5´-GACGCCCTAACACTGTGC-3´
- hush_for3: 5´-TAGTTGCCAGCCATCTGTTGT-3´
- hush_rev3: 5´-CTGAAGCCAGACAGGAAACCT-3´
- ushwt2f: 5'-CATGGTCGGCTCATGT-3'
- ushwt2r: 5'-TTGAACCTCTCGGACTATCTG-3'
- neoFOR3: 5'-GCAAGCAAGGCAGGAT-3'
- neoREV3: 5'-GTATCCCAAGGGTGGAAG-3'
- ush5arm2f: 5'-GGGTTCTGTTGGTGAG-3'
- ush5arm2r: 5'-CTAGCTTGGCCGTAAA-3'

qPCR:

- o4_qf2: 5´-CAGGTAGGTTAGGCTGATAG-3´
- o4_qr1: 5´-CAAGTTCATGAACGGCAGAAC-3´
- ng_qf6: 5´-GAGGCTTCACTTGTTAAGGG-3´
- ng_qr4: 5´-CTGAGATCAGGGTAGACATAC-3´
- ush1c_q1f: 5'-CTTCTTGGAGCAGATGGGATAAA-3'
- ush1c_q1r: 5'-GAATGCAGATTTCTGGTTTCCAC-3'

RT-PCR:
- ushrt2f: 5´-CGGAGCCTGAAGGAGCGAG-3’
- ushrt2r: 5´-TTGTTTTCCCGACTGCCAGA-3’
- urt4r: 5’-ACAGGGATCAGGCCAATGTG-3’

Gene Repair:
- ushHS1f: 5’- CCATGAAACTGACTAGTGG -3’
- huUSH2f: 5’- CCTTGCTCTGTTACCCGTTC -3’
- huUSH2r: 5’- GTTCTGTCCCAACAATCATGC -3’
- huUSH1r: 5‘- CTGGCATTTCTTGTTCTGT -3‘

(internal) sequencing primers:
- ushrt2f: 5´-CGGAGCCTGAAGGAGCGAG-3’
- ushrt2r: 5´-TTGTTTTCCCGACTGCCAGA-3’
- ushrt1s: 5´-GGTGTCAGCTGGTCGTACTCC-3’
- Ss9for: 5´- CTCAACTAATCGTGGCCTAGTG-3´
- ss11for: 5´- AGACAAGGAACTCTAATGCAAGT-3´
- ush1s: 5’- GTGCCTGGCCACATCTGGA -3’

<u>gRNAs</u> (PAM underlined, cutting site in capital letters):

rk1: GagacatattcactaacTGtg<u>GGG</u>
rk3: Gggaagtgacaggtga TCtg<u>GGG</u>
rk4: GaggccctgatcctaccAAgg<u>AGG</u>
urg1: GgacccagcacacttaCTgg<u>TGG</u>
urg2: GtaccaccagtaagtgTGct<u>GGG</u>

<u>Gene synthesis</u> (bold capital letters are added recognition sites for *Asc*I, *Not*I, *Xho*I, *Mun*I restriction enzymes; the exon 2 coding sequence is in capital letters with the nonsense causing T shaded in gray):

**GGCGCGCC**gctttgataagttaaggcagggaatttggatttgaattttgttttaagtccagtgagaatctactaagcaagcaagggacaggat
ccaatttatttaaatatttctgggccctgatcctagggaacttcagattgtttttcctgctcgaactcctccgccacaaccaaatgcaggcagctcag
gcctgtttgagtgcgattcctcacacttctcgcacaggcccaggcaaatccaggctgggtcaccatcgagatgggcttttggaggtg**GCGGCCG**
**C**gcaaca**CTCGAG**agggaaggaggtactgtcagatctaggccagaaatctgcattctgtacccccctgctcaggccagaaatcccaagggctg
ggcccagcatgtcccctctgtggtgggacggacagactgccccggtcttccagaacccttgggatacccacagaaagaggtaacgctgctctggc
cctcttctgaggacgagtcagtggagagcatgcagcttccagctgcagcctctctatgaagggctgaggccctgggccgggaggctggaggaga
gagggacccagtgacccccccaagcttccaccttgctctgttacccgttcttgggctgaagagagacccaaaaatacagtgtagagattcacactg
aggtaactcagggagtggaattcagggcctcccgctgggattgaggtgctaatgacacaactcctgaacctgaccttagagtgccagccattgac
gtcaacaaagttgaaatgatgtaacctgacgctccccctgcggggcttgtgcaggggcctggggagggggaaggagtggccatgaaactgacta
gtggacagaacccagctaaggtcaggacaagacagagtgaaggtcccctggcactgatgttacagaagaattcggtggtaaggggcttctgga
gagtggcatgtgctatctaagcgagtggcccaaatccttcctgaaagcatttatccggcactacagccaccatcaggtaagacagtgggcttcttc
tggccatggatgacacagccatgggggtgagcagcagcactgccatggcagcgtgtcactgtcacatggggattcacatatgtacctatgtgtgtt
catccccgtgtgtgcacatattgccccacctggggacaaagggtgcctggccacatctggaggggcagcggtactcctgtggccacgttggggtg
gtctgcataggtctgatgcattggggtcagaggggcagcctggcctgtggctcctcttctctcctcacaactccagccctgaaaagctgctgggga
ggcccttggggatgacctctcctccctgaggtctgctatgggggcgggtgctgagcctggagctgtgattctgctattggattttccagGTGGAT
TTTCTGATTGAAAATGATGCAGAGAAGGACTATCTCTATGATGTGCTG**<u>T</u>**GAATGTACCACCAgtaagtgtgctgggtc

cagctcttgtgggccacttgggttcctttgtcttcagggagccctgggatgggttgttctgagacagaggagctcagagggtggatgctcacggctc
ctggaaatcaaatggacataccattcactcatttcagcaactatttacacaagtactttgtacttggctttgtactaggggctgggtatagttgtgag
ccagacagattggtctctgttttcaggttgctcacagtctgatggaggaggctgtctagtagccagatagattctatagagcatgattgttgggaca
gaacaagaaatgccagctggccacagcccttgcatcagatgtctccgatcacccacttgctttttga**CAATTG**gatgggccaagagtgggcaa
gtcagctggcaggtagagaagtgccagctgcagatgttgagggtcttaatgatgtgaagctgaggctggtgggagattgtgtgagggtgggattt
gccgagctccactgcagctcagacgcaggaaccaaggggatgagagcattcaccaccctctcgcagaggtttcctgtctggcttcagaactaggc
agagctgagtttgaatcctggcttgatttcatctgcagatttccttgggatacttaaac**GGCGCGCC**

Therapeutic ssODN (with the correcting nucleotide shaded in grey and a blocking nucleotide, preventing repeated cutting by the gRNA in underlined capitals):

**urt1:** (anti-sense orientation):
5´- caacccatcccagggctccctgaagacaaaggaacccaagtggcccacaagagctggacccagcacacttactggtgAtacattc**G**cag
cacatcatagagatagtccttctctgcatcattttcaatcagaaaatccacctggaaaatccaatagcagaatcacagctccaggctcagcaccc

**urt2:** (sense-orientation):
5´- ggattttccaggtggattttctgattgaaaatgatgcagagaaggactatctctatgatgtgctg**C**gaatgtaccaccagtaagtgtgctg
gCtccagctcttgtgggccacttgggttcctttgtcttcagggagccctgggatgggttgttctgagacagaggagctcagagggtgg

**urt1.1:** (sense-orientation)
5´-tctcctccctgaggtctgctatgggtgggggtgctgagcctggagctgtgattctgctattggattttccaggtggattttctgattgaaaat
gatgcagagaaggactatctctatgatgtgctg**C**gaatgtaTcaccagtaagtgtgctgggtccagctcttgtgggccacttgggtt

**urt1.2:** (sense-orientation)
5´- atgggtgggggtgctgagcctggagctgtgattctgctattggattttccaggtggattttctgattgaaaatgatgcagagaaggactatc
tctatgatgtgctg**C**gaatgtaTcaccagtaagtgtgctgggtccagctcttgtgggccacttgggttcctttgtcttcagggagccc

**urt1.3:** (sense-orientation)
5´- tctgattgaaaatgatgcagagaaggactatctctatgatgtgctg**C**gaatgtaTcaccagtaagtgtgctgggtccagctcttgtgggc
cacttgggttcctttgtcttcagggagccctgggatgggttgttctgagacagaggagctcagagggtggatgctcacggctcctggaaa

**urt1.4:** (anti-sense-orientation)
5´- aacccaagtggcccacaagagctggacccagcacacttactggtgAtacattc**G**cagcacatcatagagatagtccttctctgcatcatt
ttcaatcagaaaatccacctggaaaatccaatagcagaatcacagctccaggctcagcacccccacccatagcagacctcagggaggaga

**urt1.5:** (anti-sense-orientation)
5´- tttccaggagccgtgagcatccaccctctgagctcctctgtctcagaacaacccatcccagggctccctgaagacaaaggaacccaagtg
gcccacaagagctggacccagcacacttactggtgAtacattc**G**cagcacatcatagagatagtccttctctgcatcattttcaatcaga

### References


Albl, B., Haesner, S., Braun-Reichhart, C., Streckel, E., Renner, S., Seeliger, F., Wolf, E., Wanke, R., and Blutke, A. (2016). Tissue Sampling Guides for Porcine Biomedical Models. Toxicol Pathol *44*, 414-420.

Andersson, R., Gebhard, C., Miguel-Escalada, I., Hoof, I., Bornholdt, J., Boyd, M., Chen, Y., Zhao, X., Schmidl, C., Suzuki, T*., et al.* (2014). An atlas of active enhancers across human cell types and tissues. Nature *507*, 455-461.

Barone, F., Nannoni, E., Elmi, A., Lambertini, C., Scorpio, D.G., Ventrella, D., Vitali, M., Maya-Vetencourt, J.F., Martelli, G., and Benfenati, F. (2018). Behavioral Assessment of Vision in Pigs. Journal of the American Association for Labaratory Animal Science *57*, 350-356.

Brudno, M., Steinkamp, R., and Morgenstern, B. (2004). The CHAOS/DIALIGN WWW server for multiple alignment of genomic sequences. Nucleic Acids Res *32*, W41-44.

Ferretti, V., Poitras, C., Bergeron, D., Coulombe, B., Robert, F., and Blanchette, M. (2007). PReMod: a database of genome-wide mammalian cis-regulatory module predictions. Nucleic Acids Res *35*, D122-126.

Fishilevich, S., Nudel, R., Rappaport, N., Hadar, R., Plaschkes, I., Iny Stein, T., Rosen, N., Kohn, A., Twik, M., Safran, M*., et al.* (2017). GeneHancer: genome-wide integration of enhancers and target genes in GeneCards. Database (Oxford) *2017*.

Giresi, P.G., Kim, J., McDaniell, R.M., Iyer, V.R., and Lieb, J.D. (2007). FAIRE (Formaldehyde-Assisted Isolation of Regulatory Elements) isolates active regulatory elements from human chromatin. Genome Res *17*, 877-885.
Hood, D.C., Bach, M., Brigell, M., Keating, D., Kondo, M., Lyons, J.S., Marmor, M.F., McCulloch, D.L., Palmowski-Wolfe, A.M., and International Society For Clinical Electrophysiology of, V. (2012). ISCEV standard for clinical multifocal electroretinography (mfERG) (2011 edition). Doc Ophthalmol *124*, 1-13.
Karlstetter, M., Sorusch, N., Caramoy, A., Dannhausen, K., Aslanidis, A., Fauser, S., Boesl, M.R., Nagel-Wolfrum, K., Tamm, E.R., Jägle, H., *et al.* (2014). Disruption of the retinitis pigmentosa 28 gene Fam161a in mice affects photoreceptor ciliary structure and leads to progressive retinal degeneration. Hum Mol Genet *23*, 5197-5210.
Klymiuk, N., Mundhenk, L., Kraehe, K., Wuensch, A., Plog, S., Emrich, D., Langenmayer, M.C., Stehr, M., Holzinger, A., Kroner, C., *et al.* (2012). Sequential targeting of CFTR by BAC vectors generates a novel pig model of cystic fibrosis. J Mol Med (Berl) *90*, 597-608.
Kooijman, A.C., and Damhof, A. (1986). A tricolor light source for stimulation and adaptation in electroretinography. Doc Ophthalmol *63*, 195-203.
Kurome, M., Kessler, B., Wuensch, A., Nagashima, H., and Wolf, E. (2015). Nuclear transfer and transgenesis in the pig. Methods Mol Biol *1222*, 37-59.
Luo, J., Ying, K., and Bai, J. (2005). Savitzky–Golay smoothing and differentiation filter for even number data. Signal Processing *85*, 1429-1434.
McLean, D.J., and Skowron Volponi, M.A. (2018). trajr: An R package for characterisation of animal trajectories. Ethology *124*, 440-448.
Overlack, N., Kilic, D., Bauss, K., Marker, T., Kremer, H., van Wijk, E., and Wolfrum, U. (2011). Direct interaction of the Usher syndrome 1G protein SANS and myomegalin in the retina. Biochim Biophys Acta *1813*, 1883-1892.
Reiners, J., Reidel, B., El-Amraoui, A., Boeda, B., Huber, I., Petit, C., and Wolfrum, U. (2003). Differential distribution of harmonin isoforms and their possible role in Usher-1 protein complexes in mammalian photoreceptor cells. Invest Ophthalmol Vis Sci *44*, 5006-5015.
Richter, A., Kurome, M., Kessler, B., Zakhartchenko, V., Klymiuk, N., Nagashima, H., Wolf, E., and Wuensch, A. (2012). Potential of primary kidney cells for somatic cell nuclear transfer mediated transgenesis in pig. BMC Biotechnol *12*, 84.
Savitzky, A., and Golay, M.J.E. (1964). Smoothing and Differentiation of Data by Simplified Least Squares Procedures. Analytical Chemistry *36*, 1627-1639.
Schindelin, J., Rueden, C.T., Hiner, M.C., and Eliceiri, K.W. (2015). The ImageJ ecosystem: An open platform for biomedical image analysis. Mol Reprod Dev *82*, 518-529.
Sedmak, T., and Wolfrum, U. (2010). Intraflagellar transport molecules in ciliary and nonciliary cells of the retina. J Cell Biol *189*, 171-186.
Sedmak, T., and Wolfrum, U. (2011). Intraflagellar transport proteins in ciliogenesis of photoreceptor cells. Biol Cell *103*, 449-466.
Sorusch, N., Bauss, K., Plutniok, J., Samanta, A., Knapp, B., Nagel-Wolfrum, K., and Wolfrum, U. (2017). Characterization of the ternary Usher syndrome SANS/ush2a/whirlin protein complex. Hum Mol Genet *26*, 1157-1172.
Thurman, R.E., Rynes, E., Humbert, R., Vierstra, J., Maurano, M.T., Haugen, E., Sheffield, N.C., Stergachis, A.B., Wang, H., Vernot, B., *et al.* (2012). The accessible chromatin landscape of the human genome. Nature *489*, 75-82.
Trojan, P., Krauss, N., Choe, H.W., Giessl, A., Pulvermuller, A., and Wolfrum, U. (2008). Centrins in retinal photoreceptor cells: regulators in the connecting cilium. Prog Retin Eye Res *27*, 237-259.
van Wijk, E., van der Zwaag, B., Peters, T., Zimmermann, U., Te Brinke, H., Kersten, F.F., Marker, T., Aller, E., Hoefsloot, L.H., Cremers, C.W., *et al.* (2006). The DFNB31 gene product whirlin connects to the Usher protein network in the cochlea and retina by direct association with USH2A and VLGR1. Hum Mol Genet *15*, 751-765.
Vochozkova, P., Simmet, K., Jemiller, E.M., Wunsch, A., and Klymiuk, N. (2019). Gene Editing in Primary Cells of Cattle and Pig. Methods Mol Biol *1961*, 271-289.

Wang, J., Zhuang, J., Iyer, S., Lin, X.Y., Greven, M.C., Kim, B.H., Moore, J., Pierce, B.G., Dong, X., Virgil, D., *et al.* (2013). Factorbook.org: a Wiki-based database for transcription factor-binding data generated by the ENCODE consortium. Nucleic Acids Res *41*, D171-176.
Warming, S., Costantino, N., Court, D.L., Jenkins, N.A., and Copeland, N.G. (2005). Simple and highly efficient BAC recombineering using galK selection. Nucleic Acids Res *33*, e36.
Waterhouse, A.M., Procter, J.B., Martin, D.M., Clamp, M., and Barton, G.J. (2009). Jalview Version 2--a multiple sequence alignment editor and analysis workbench. Bioinformatics *25*, 1189-1191.
